# Hepatitis B virus HBx protein masks epigenetic reader Spindlin1 via an inter-molecular zinc finger to subvert transcriptional control

**DOI:** 10.1101/2025.11.05.686744

**Authors:** Alexis Clavier, Toshinobu Shida, Santiago Gómez-Evain, Maxim A. Droemer, Franziska von Hammerstein, Julian Holzinger, Mila M. Leuthold, Anne K. Schütz

## Abstract

The HBx protein from Hepatitis B virus (HBV) is essential for replication and promotes pathogenicity during chronic infection. HBx hijacks host proteins, reprogramming them to evade antiviral defence. However, the structural basis of recruitment remains largely unknown. The HBx_1-120_ isoform derives from integration of viral DNA into the host genome and is linked to hepatocellular carcinoma.

We present an NMR-based setup to characterize HBx-host protein interactions at residue-level resolution. HBx_1–120_ is disordered in isolation but undergoes local folding upon binding apoptosis regulator Bcl-xL and epigenetic reader Spindlin1. The HBx-Spindlin1 complex is bivalent: a hydrophobic interaction combines with a sticky patch realized via a rare inter-molecular zinc finger. HBx thus conceals the region responsible for recruiting Spindlin1 to histone tails, promoting transcription of extrachromosomal viral DNA. This mechanism exemplifies the capacity of HBx to engage diverse host factors into dynamic complexes.

## Introduction

The Hepatitis B virus (HBV) remains a global health challenge. Chronic HBV carriers, whose immune system fails to eradicate the virus, are at high risk of developing liver cirrhosis and hepatocellular carcinoma (HCC).^1^ The HBV genome encodes four proteins. Structural biology has focused on the components that make up the virion^2,3^: the prototypical capsid shell that hosts the viral genome and the polymerase^4,5^, the membrane proteins in the lipid envelope^6^, and receptor binding during viral uptake^7^. In infected hepatocytes, the viral DNA persists in a chromatinized form termed cccDNA (covalently closed circular)^8^, which represents the central reservoir for viral chronicity.

Besides the structural components, HBV encodes the regulatory protein X (HBx)^9^ (Fig. 1A); its structure and functions remain poorly understood. The 154-residue protein bears little homology to other proteins and is essential not to infection but to replication^10^. While mammalian *hepadnaviruses* encode an X gene, which is highly conserved among apes (Fig. S1A), their avian counterparts lack HBx, which correlates with lack of chronic hepatitis and HCC in these natural hosts^11,12^. The expression of HBx is sufficient to induce HCC in mouse models^13,14^. A link exists between HCC and the presence of the short isoform HBx_1-120_ found in many chronically infected patients^14,15^. Transcripts of this isoform derive from viral DNA that integrates into the host genome at the site of double-strand breaks and persists independently of the cccDNA. Although the mechanism of carcinogenesis remains unclear, HBx is regarded as one of the most important – if not the primary – oncogenes of HBV.

**Figure 1.**
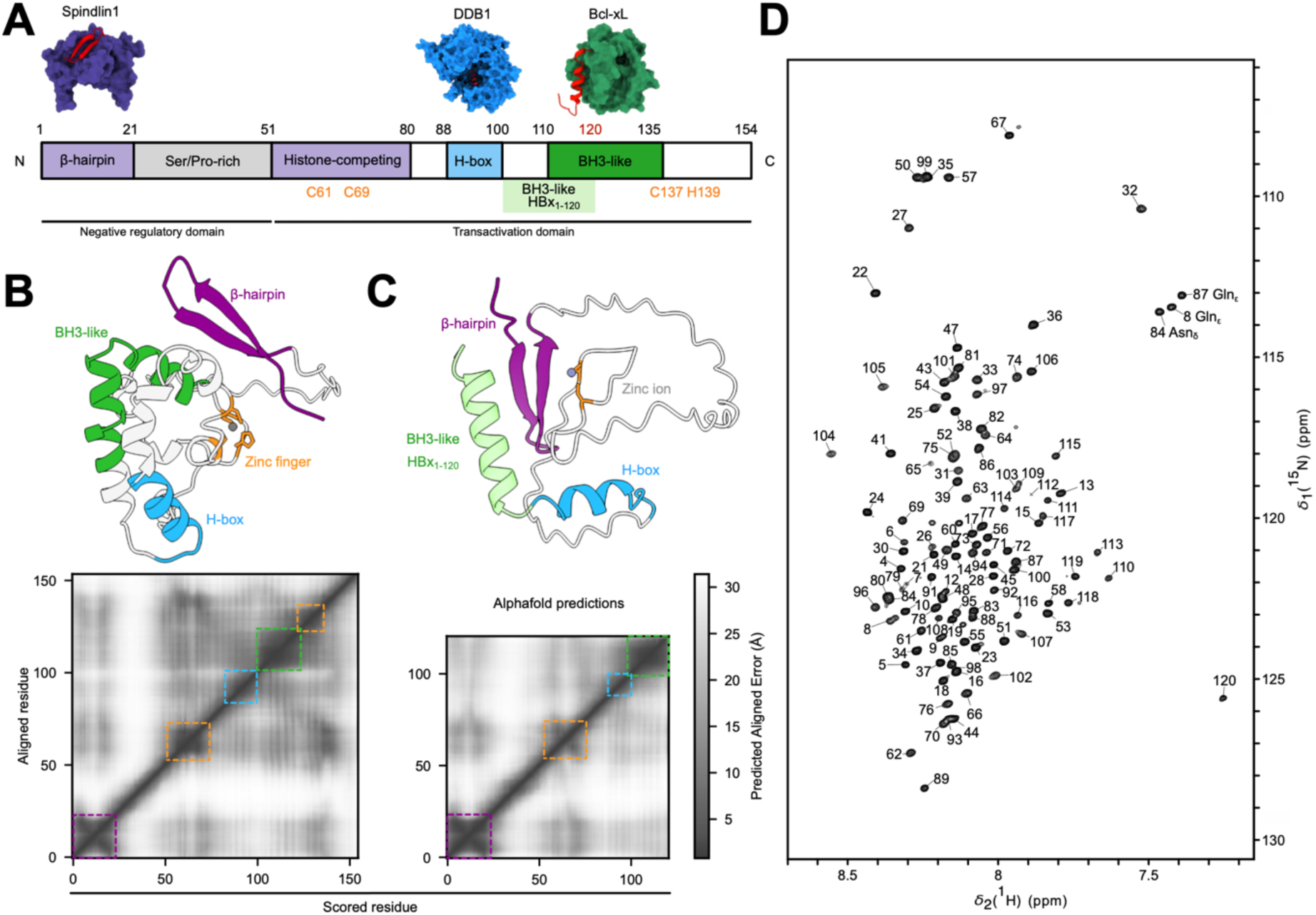
Structural elements and disorder in HBx isoforms. **A.** Schematic HBx protein sequence with colours highlighting structured elements observed in complex with host proteins and regions with defined characteristics. Highly conserved residues suspected to form an intra-molecular zinc finger based on mutagenesis experiments^17^ are indicated. **B.** AlphaFold (AF)^30^ model for full-length HBx_1-154_ with a Zn^2+^ ion. The β-hairpin binding Spindlin1, two helices (H-box, BH3-like) that recognize DDB1 and Bcl-2/xL, respectively, and the putative zinc finger are highlighted. However, the PAE (predicted aligned error) matrix indicates that the overall fold is poorly defined, consistent with HBx displaying intrinsic disorder (*cf*. parameters in Tab. S3). **C.** AF model for isoform HBx_1-120_ with a Zn^2+^ ion. Compared to HBx_1-154_, the BH3-like helix is shortened and shifted toward the N-terminus^31^; the putative intra-molecular zinc finger is lost. The PAE map reveals lower confidence in the overall topology. **D.** Amide correlation spectrum (SOFAST ^1^H-^15^N HMQC, here and all subsequent figures) of HBx_1-120_ in the presence of residual urea (190 mM urea, 125 mM L-arginine at 4°C). The low ^1^H chemical shift dispersion indicates intrinsic disorder. Amide and sidechain assignments were obtained except for three invisible residues at the N-terminus^27^.

HBx is the earliest HBV protein expressed post infection^16^ and predominantly localizes to the nucleus^17^. It disables several host restriction factors and thereby subverts the host cell’s antiviral defence and creates a cellular environment that supports viral replication. Nuclear HBx activates transcription from the extrachromosomal cccDNA by hijacking chromatin-associated host factors to overcome cellular repression. This process notably involves recruitment of DNA damage-binding protein 1 (DDB1), a core component of ubiquitin ligase complexes^18^, and the chromatin reader Spindlin1^19^. In the cytoplasm, HBx interferes with signalling pathways, including mitochondrial apoptosis regulation via Bcl-family proteins^20^. HBx is a multifunctional protein with over 200 proposed interactors, most of which are supported only by pull-down assays and lack further validation^21^. Crystal structures of a few host proteins bound to short HBx- derived peptides^22–25^ reveal the recognition modes of short linear motifs within HBx. However, the structural mechanisms by which HBx diverts host protein functions remain unknown, due to challenges in reconstituting native HBx *in vitro*. A cryo-electron microscopy structure of a complex with DDB1 attained low local resolution of HBx^25^, hinting at a dynamic state and reinforcing the notion of HBx being an intrinsically disordered protein (IDP).

Here, we investigate the isoform HBx_1-120_ using NMR spectroscopy. In complex with two host proteins potentially implicated in carcinogenesis, Bcl-xL and Spindlin1, HBx_1-120_ remains predominantly dynamically disordered and folds only locally. HBx_1-120_ wraps around Spindlin1, driven by a short interaction motif that mimics human transcriptional regulator SPINDOC, in combination with an inter-molecular zinc finger, a rare motif in virus-host protein interactions. HBx thus competes with methylated histone tails, the physiological interaction partners of Spindlin1, thereby interfering with transcription regulation.

Our NMR-based approach overcomes the challenges of solving a putative HBx ‘structure’ and establishes an experimental framework for residue-level interaction studies of a large HBx segment. Moreover, we reveal a previously undocumented principle by which a viral IDP combines minimal elements within its short sequence to accommodate diverse host protein interactions.

### HBx_1-120_ is dynamically disordered in the absence of binding partners

AlphaFold (AF) predictions of full-length HBx_1-154_ in the presence of a Zn^2+^ ion recapitulate the short structural motifs known from crystal structures and the putative zinc finger^17,22–24^ (Fig. 1A-B). All reported secondary structure elements are accommodated simultaneously, albeit with low confidence, particularly for long-range contacts. Comparing full-length HBx_1-154_ and the HCC-associated isoform HBx_1-120_, the truncation is expected to change host protein interactions, considering that two of the residues from the intra-molecular zinc finger are absent. Indeed, AF predicts an overall less compact fold at lower confidence (Fig. 1C).

Experimental verification of AF predictions requires integral HBx, which has so far resisted crystallization, high-resolution cryo-EM and NMR studies. Meanwhile, we successfully stabilized the isoform HBx_1-120_ *in vitro* without solubilization tags or modifications^26^ and obtained the sequential backbone NMR resonance assignments^27^ (Fig. 1D). NMR samples of HBx_1-120_ can be prepared at concentrations up to 50–100 𝜇M and remain stable for 24-36 hours. While relaxation and dynamics experiments are limited by low sensitivity, this setup enables fingerprinting of amide and methyl groups to probe conformational dynamics and interactions. The amide correlation spectrum of HBx_1-120_ in isolation (Fig.1D) reveals low dispersion in the ^1^H dimension, indicating a dynamically disordered character in the presence of residual urea (190 mM). Secondary structure prediction from ^13^C backbone shifts, which could only be obtained in the presence of 1 M urea, corroborate a lack of stable secondary structure except for some propensity for a BH3-like helix in the C-terminal region (Fig. S1B). The translational diffusion coefficient *D* is inversely related to the hydrodynamic radius and reports on the compaction of a polypeptide chain. The *D* value of HBx_1-120_ derived from DOSY-NMR experiments contradicts the compacted, folded state of HBx_1-120_ predicted by AF (Tab. 1). Taken together, the experimental data suggest that HBx_1-120_ is predominantly disordered in the absence of binding partners.

**Table 1.** Analysis of diffusion coefficients as proxy for molecular compaction. Comparison of experimental translational diffusion coefficient *D* obtained by DOSY-NMR with predictions based on structure (HYDROPRO^32^ predictions-based crystal structures or AF models) and with empirical estimations (cf. Methods and Tab. S2)^33^. The experimental NMR setup combines selective methyl labelling of HBx_1-120_ with a ^13^C filter to ensure that diffusion is measured of HBx_1-120_ only, with no contribution of the binding partners. Despite its lower molecular weight, HBx_1-120_ diffuses more slowly than Bcl-xL and Spindlin1. The *D* coefficient measured for HBx_1-120_ lies between the values predicted from the AF model and those expected for a pure IDP. In the presence of Spindlin1, HBx_1-120_ diffuses more slowly, indicating binding without compaction into a tight complex. If HBx wrapped tightly around Spindlin1, the *D* coefficient would approach that of Spindlin1 alone. Indeed, addition of Zn^2+^ causes an increase in diffusion, consistent with compaction into a tighter complex. The diffusion of HBx_1-120_ is not significantly affected by the presence of Bcl-xL. At protein concentrations of 50 𝜇M used for DOSY experiments, the low affinity (∼200 𝜇M, Tab. S4) of the modified BH3-like helix in HBx_1-120_ (Fig. 2C) precludes the formation of tight binary or ternary complexes, even if transient interactions are detectable in NMR fingerprint spectra (Fig. 2).

| Protein or complex | Molecular weight [kDa] | *D* experimental [10<sup>-11</sup> m<sup>2</sup>/s] | *D* predicted from structure [10<sup>-11</sup> m<sup>2</sup>/s] | *D* predicted empirical [10<sup>-11</sup> m<sup>2</sup>/s] |
| --- | --- | --- | --- | --- |
| Bcl-xL | 19.99 | 4.44 +/- 0.06 | 4.68 | 4.43 (folded) |
| Spindlin1 | 24.7 | 4.97 +/- 0.12 | 4.12 | 4.1 (folded) |
| HBx<sub>1-120</sub> | 12.85 | 3.74 +/- 0.08 | 4.00 | 3.41 (IDP) |
| HBx<sub>1-120</sub> - Spindlin1 | 37.55 | 3.32 +/- 0.09 | 3.57 |  |
| HBx<sub>1-120</sub> - Spindlin1 - Zn<sup>2+</sup> | 37.55 | 4.35 +/- 0.08 | 3.42 |  |
| HBx<sub>1-120</sub> - Bcl-xL | 32.84 | 3.81 +/- 0.08 | 3.19 |  |
| HBx<sub>1-120</sub> - Bcl-xL - Spindlin1 | 57.54 | 3.24 +/- 0.03 | 3.04 |  |

Viral IDPs can hijack host proteins through diverse recognition modes, including coupled folding and binding – either globally or locally at one or more interaction motifs – as well as electrostatically driven and highly dynamic complexes^28,29^. We investigated the molecular mechanisms by which HBx_1-120_ engages two host proteins, Bcl-xL and Spindlin1.

### HBx_1-120_ interacts transiently with apoptosis regulator Bcl-xL

We benchmarked our NMR setup for HBx_1-120_ using an established binding partner, the soluble domain of Bcl-xL. The HBx_113-135_ peptide forms a BH3-like helix that co-crystallises with Bcl-xL^22,34,35^; peptides derived from the HBx_1-120_ isoform, in which the BH3-like helix is partially truncated, show a marked reduction in affinity^31^ (Fig. 1A).

Complex formation between HBx_1-120_ and Bcl-xL could be observed in several NMR experiments. The interaction manifests in complete intensity loss in amide correlation spectra in the C-terminal region of HBx_1-120_ (Fig. 2A). The interaction is monovalent and localized strictly to the shifted BH3-like helix covering residues 100–120, in line with the AF prediction (Fig. 2B). As helical propensity in this region already exists for HBx_1-120_ in isolation (Fig. S1B), this interaction appears to be driven by a conformational selection mechanism. In methyl-detected spectra of HBx_1-120_ alone, the signals of Ile, Leu, Val and Met residues appear at the random coil positions. Upon addition of Bcl-xL, signals from three Leu and one Met residues inside the BH3-like helix disappear (Fig. 2C). The interaction is likewise reflected in signals of methyl groups on Bcl-xL that surround its BH3 binding groove (Fig. 2D). Again, the signals show mild signal attenuation without chemical shift perturbations (CSPs). The signal loss is attributed to conformational exchange on the ms–𝜇s time scale due to transient interface formation. Methyl groups involved in a stable hydrophobic interface would remain detectable within the complex, thanks to the methyl-TROSY effect.

**Figure 2.**
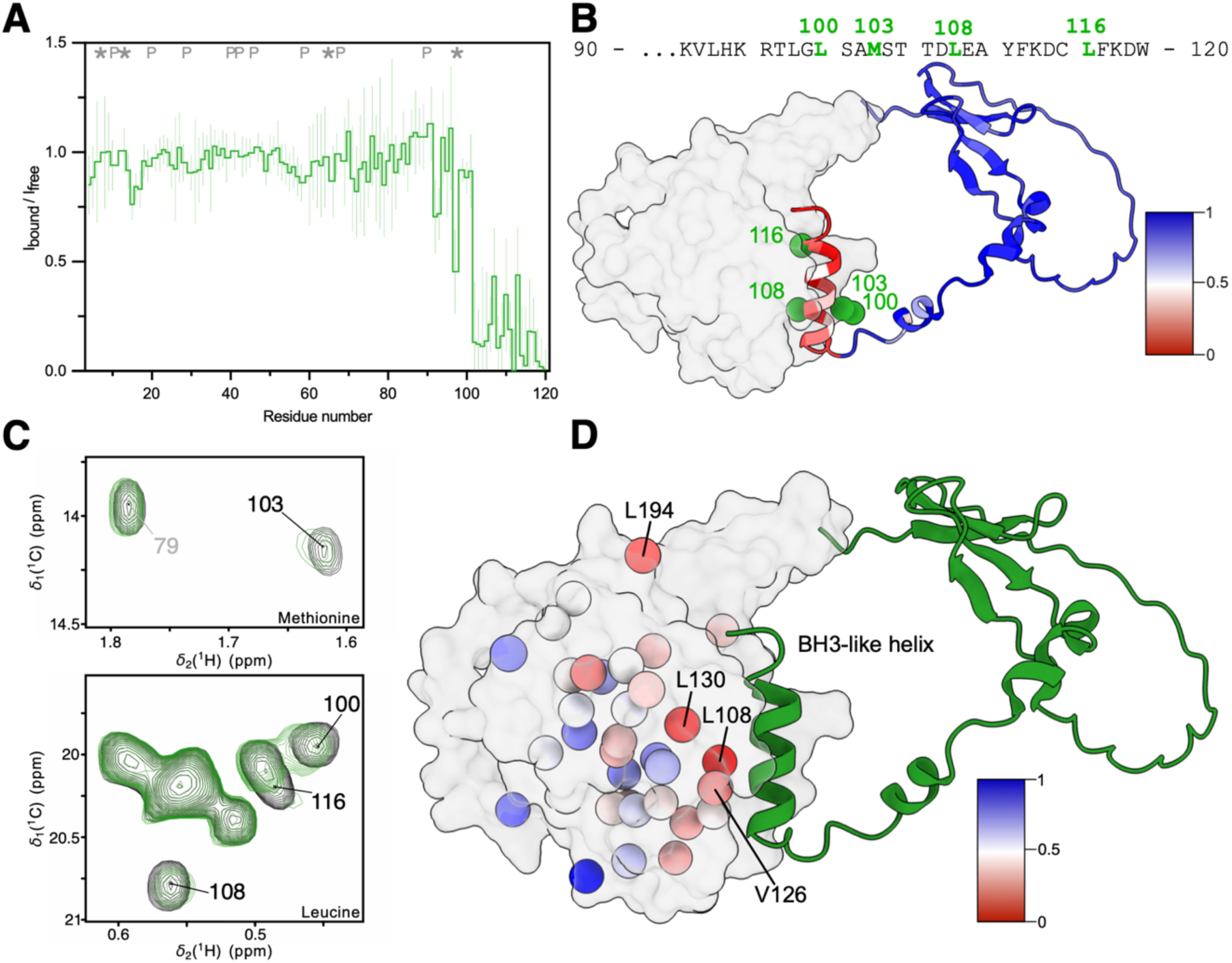
The interaction between HBx_1-120_ and Bcl-xL is monovalent. **A.** NMR signals of HBx_1-120_ exhibit intensity loss for residues 100–120 upon complex formation with Bcl-xL (1:1 stochiometric ratio, 50 µM); ‘P’ indicates proline residues, which lack observable amide correlations; ‘*’ marks residues excluded due to spectral overlap precluding peak-height estimations. **B.** Intensity loss is visualized as the intensity ratio I_bound_/I_free_ (color-coded from red to blue, representing maximal to minimal signal attenuation) on the AF model of the complex of Bcl-xL (surface representation) and HBx_1-120_ (cartoon). The predicted and NMR-derived (red) interacting regions are identical. Signal attenuation arises from an increase in transverse relaxation rates as the BH3-like helix associates with Bcl-xL, effectively more than tripling the complex size, with likely contributions from chemical-exchange broadening. AF predicts the N-terminal part of HBx_1-120_ to form a β-hairpin, for which there is no experimental evidence (PAE matrix in Fig. S2B). **C.** Comparison of free (black contours) and Bcl-xL bound (green) methyl-group correlations of HBx_1-120_ (ILVM-^13^CH_3_-labelling) in ^1^H-^13^C SOFAST HMQC spectra reveals signal loss for four methyl probes in the BH3-like helix region of HBx_1-120_ (black labels), with residues outside this region unaffected (grey label). **D.** Bcl-xL-detected experiments reveal specific intensity loss for methyl groups surrounding the BH3-binding groove in the presence of HBx_1-120_. The AF model (same as in panel B) is coloured according to the normalized intensity ratio I_bound_/I_free_ (see details in Fig. S2D, E).

The transient nature of the complex is further corroborated by DOSY-NMR (Tab. 1). The diffusion coefficient *D* reflects on two competing processes that modulate the hydrodynamic radius of HBx_1-120_: an increase in molecular size from complex formation reduces *D*, while compaction of the disordered HBx_1-120_ due to folding increases *D*. The diffusion coefficients of HBx_1-120_ alone and in the presence of Bcl-xL are identical, suggesting that both proteins do not effectively diffuse together. This is consistent with binding affinities on the order of hundreds of 𝜇M^31^. The interaction is restricted to the C-terminal helix and does not induce compaction of HBx_1-120_ through folding. The modulation of apoptosis by HBx is context-dependent^36,37^. Our analysis confirms that the HBx_1-120_ isoform is, at best, a weak Bcl-xL interactor, incapable of competing with host proteins vying for the same binding site^31,38^. Therefore, this interaction is unlikely to be responsible for the carcinogenicity of the short isoform.

In sum, our NMR setup is sufficiently sensitive to characterize even transient, low-affinity interactions of HBx_1-120_ at the residue level, such as the monovalent binding mode of Bcl-xL.

### HBx_1-120_ engages Spindlin1 in a bivalent manner

The epigenetic reader Spindlin1 regulates transcription and features three distinct interaction sites across its three Tudor domains (Fig. 3A)^39–41^. The first two sites recognize methylated histone tails, while the third acts as a docking site for the regulatory protein SPINDOC (C11orf84; Fig. 5B)^42^. A peptide HBx_2-21_ derived from the N-terminal region of HBx forms a β-hairpin that completes the third Tudor β-barrel of Spindlin1^24^. We also observe this interaction in NMR experiments with integral HBx_1-120_, manifested as signal loss in amide-detected spectra (Fig. 3B,C) and as CSPs for residues near the exit of the binding region (Fig. 3D). Besides that, less pronounced intensity losses are detected in the region around residues 50–80, which suggests a second, possibly weaker, interaction region, previously hypothesized from mutagenesis experiments^19^. Secondary chemical shifts (Fig. S1) indicate lack of secondary structure in the absence of Spindlin1, consistent with a folding upon binding mechanism. A double mutation in the third Tudor domain of Spindlin1, designed to break the hydrophobic interface with the HBx β-hairpin, abolishes the interaction, identifying this interface as the primary driving force (Fig. 3B). We obtained an AF model of the Spindlin1-HBx_1-120_ complex (Fig. 3C) predicting two interaction regions: the β-hairpin and a second interface spanning the first two Tudor domains of Spindlin1 and residues 50–80 of HBx_1-120_. Remarkably, the intensity loss in NMR maps onto the exact interaction sites identified in the prediction. We recorded methyl-detected spectra of HBx_1-120_ that report on hydrophobic contacts in interface regions with Spindlin1. Upon binding, folding-induced CSPs move signals away from the random coil values to isolated positions. This effect was observed for two Val and four Leu signals, consistent with their presence in the β-hairpin region of HBx_1-120_ (Fig. 3F). In contrast to Bcl-xL, binding of HBx_1-120_ to Spindlin1 caused a reduction in the diffusion coefficient *D* determined from DOSY NMR, suggesting association of the two proteins into a complex with an increased hydrodynamic radius (Tab. 1).

**Figure 3.**
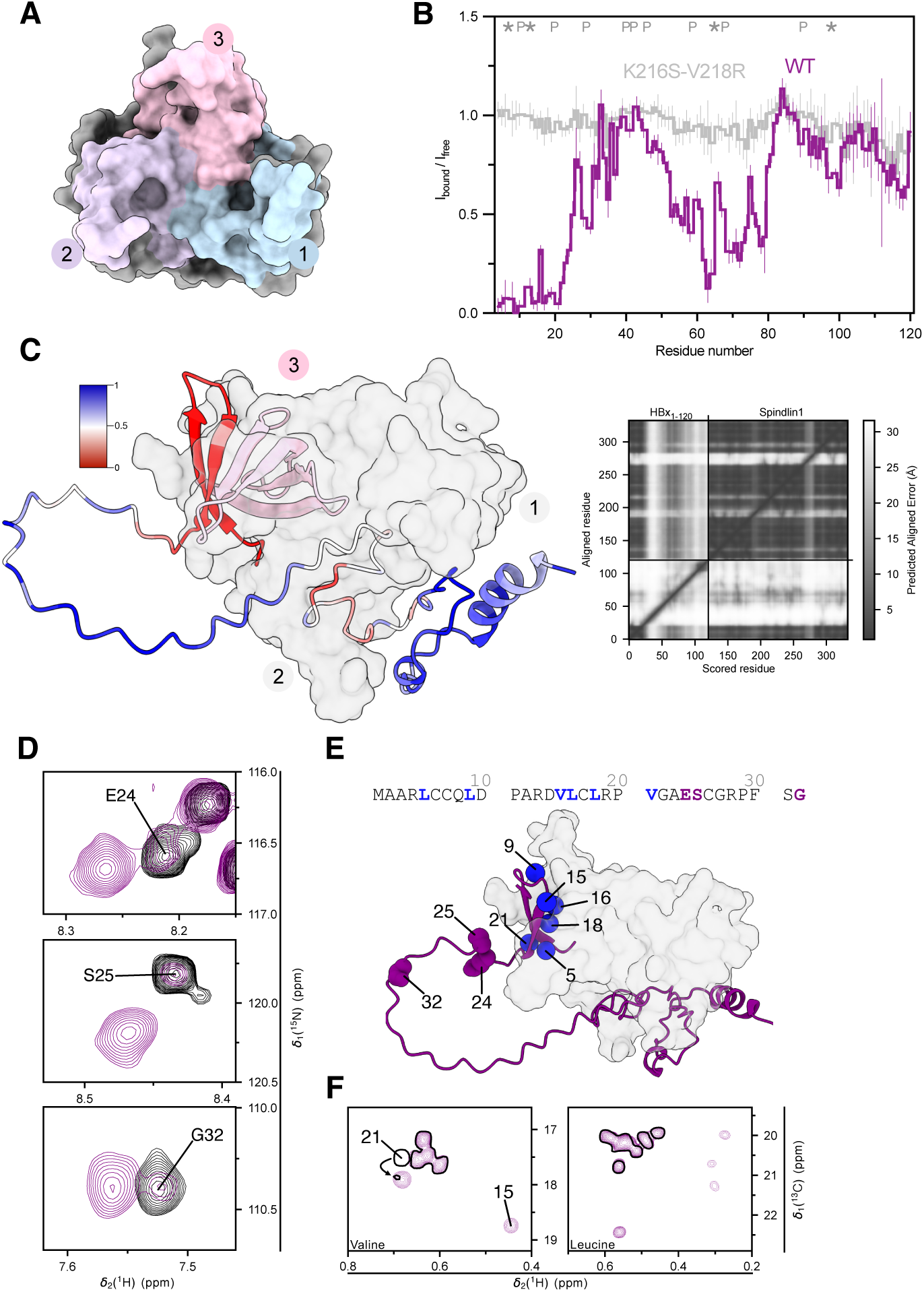
The interaction between HBx_1-120_ and Spindlin1 is bivalent. **A.** Spindlin1 is composed of three Tudor domains each forming a β-barrel: Tudor 1 (residues 54-103) recognizes methylation at K9 of histone H3; Tudor 2 (residues 133-182) recognizes methylation at K4 of H3; Tudor 3 (residues 214-259) features an incomplete β-barrel for docking of transcriptional regulators via a β-hairpin (e.g. SPINDOC in Fig. S7C or HBx residues 1–30 in panel C). **B.** NMR interaction analysis, expressed as the intensity ratio I_bound_/I_free_ for amide signals in free HBx_1-120_ and in complex with Spindlin1 (50 µM, 1:1 ratio). Signal attenuation, indicative of binding, is observed for HBx_1-120_ residues in regions 1–30 and 50–80 (full spectra in Fig. S3A). A Spindlin1 double mutation^24^ disturbing the interaction with the HBx β-hairpin causes a complete loss of binding also for residues 50–80. ‘P’ indicates proline residues; ‘*’ residues excluded due to signal overlap. **C**. AF model of the Spindlin1-HBx_1-120_ complex (in surface and cartoon representation, respectively), coloured according to intensity ratios from panel B, showing strong agreement between experimental and predicted interaction regions. Residues 50–80 of HBx_1-120_ engage the first two Tudor domains of Spindlin1, which normally recognize the methylated H3 histone tail. The PAE map indicates higher confidence of the position of the β-hairpin compared to the second binding region. **D.** Excerpts from amide correlation spectra of free HBx_1-120_ (black) versus Spindlin1-bound (purple). Residues in the flexible loop connecting the two binding regions (panel E) display chemical shift perturbations (CSPs), indicating subtle changes in backbone conformation upon complex formation. **E.** Same AF model as in panel C, with HBx_1-120_ residues of interest highlighted in purple and blue to indicate amide and methyl group CSPs, respectively. **F.** In methyl correlation spectra of free (black) and Spindlin1-bound HBx_1-120_ (purple), CSPs are observed for residues located at the hydrophobic interface of the β-hairpin. Four peaks in the Leu region appear far from the random coil position in the spectrum of the complex, consistent with four Leu residues (5, 9, 16, 18) in the β-hairpin.

Functionally, HBx can bridge two host proteins into a ternary complex that would normally not associate in its absence. Specifically, HBx engages the structural maintenance of chromosomes complex (SMC5/6), which acts as a restriction factor that inhibits transcription from the cccDNA, and bridges it to a DDB1-containing E3 ubiquitin ligase complex, thereby targeting this HBV antagonist for degradation^18^. Similarly, ternary complex formation with DDB1 and Spindlin1 has been reported^25^, although the biological function remains elusive. On the molecular level, NMR results support the concept that HBx_1-120_ can engage Bcl-xL and Spindlin1 simultaneously to form a ternary complex. Bridging between N- and C-terminal binding partners is evident from amide correlation intensity analysis (Fig. S3D). There is no evidence for cooperativity within this complex; the signal attenuations observed for the individual interaction motifs in isolation simply combine. Given that Bcl-xL localizes to the mitochondrial membrane and Spindlin1 to the nucleus, the complex is unlikely to be biologically relevant. Nevertheless, the experiment confirms that N- and C-terminal interactions of HBx_1-120_ are not mutually exclusive, provided the local recognition motifs do not overlap.

### HBx_1–120_ binds Spindlin1 in a zinc-dependent manner

The second Spindlin1 interaction region in HBx_1-120_ (residues 50–80) displays weaker signal attenuation (Fig. 3B) than the first region (residues 1–30). The complex is not fully defined in the AF prediction, as indicated by the predicted alignment error (PAE, Fig. 3C) and the observation of two distinct binding geometries (Fig. S3C). The diffusion coefficient *D* suggests a loosely associated complex (Tab. 1). These observations led us to hypothesize that a critical component is missing to form a tight, high-affinity complex in our *in vitro* setup.

The HBx_1-120_ isoform lacks two of the cysteine residues forming the putative intra-molecular zinc-finger motif (Fig. 1A). Still, dialysis of Zn^2+^ ions into the HBx_1-120_-Spindlin1 complex causes an overall loss of amide signal intensity, most pronounced for residues 35-50 (Fig. S4A-B), and this effect is reversible upon Zn^2+^ chelation (Fig. S4A-B). The signal loss is accompanied by a pronounced increase in the diffusion coefficient, approaching that of Spindlin1 alone, indicating that HBx_1-120_ compacts as it wraps more tightly around Spindlin1 (Tab. 1). When AF was provided with a single Zn^2+^ ion, it predicted an inter-molecular zinc finger to which Spindlin1 contributes C96 and H252, and HBx_1-120_ contributes C61 and C69. Two topologies of HBx_1-120_ were suggested, encompassing the Tudor domains in either the 3-2-1 or 3-1-2 order (Fig. S4D). Compared to the AF models without Zn^2+^ (Fig. S3C), the confidence in the PAE map increases slightly for the second binding region that forms the zinc finger. It is common for IDPs to sample several bound topologies in a dynamic equilibrium, with only one out of multiple linear recognition motifs (here the β-hairpin) adopting a fixed structure^43^. The most intense amide signals remaining detectable in the Zn^2+^-bound complex originate from the extended linker connecting the two interaction sites, which consists of a Ser/Pro-rich region (Fig. 1A). CSPs observed in the linker region upon binding Spindlin1 alone are enhanced upon Zn^2+^ addition, which also induces new CSPs (Fig. 4C, Fig. S4). Methyl-detected spectra of HBx_1-120_ indicate subtle changes to hydrophobic contacts in the presence of Zn^2+^. The signals of the residues closest to C61—V60 and L58—disappear from their random coil position (Fig. 4D). We performed a control experiment with HBx_1-120_ and Spindlin1 variants in which the zinc finger residues CCCH are mutated to AAAF. These mutations do not abolish the binding of HBx_1-120_ to Spindlin1 but eliminate its sensitivity to Zn^2+^ ions (Fig. S5). To complement the validation of the structural model, we performed Spindlin1-detected experiments. High-resolution NMR spectra of this folded protein require temperatures above 4°C, which are not compatible with HBx_1-120_ due to aggregation. We therefore reduced HBx to the minimal region required to bind Tudor domains 1 and 2 (HBx_56–76_; Fig. 5B). Binding of this peptide induces CSPs in the amide correlation spectrum of Spindlin1, which are enhanced in the presence of Zn^2+^ (Fig. S9C). This is consistent with the observations above that HBx_1-120_ and Spindlin1 can form a so far undescribed inter-molecular zinc finger.

**Figure 4.**
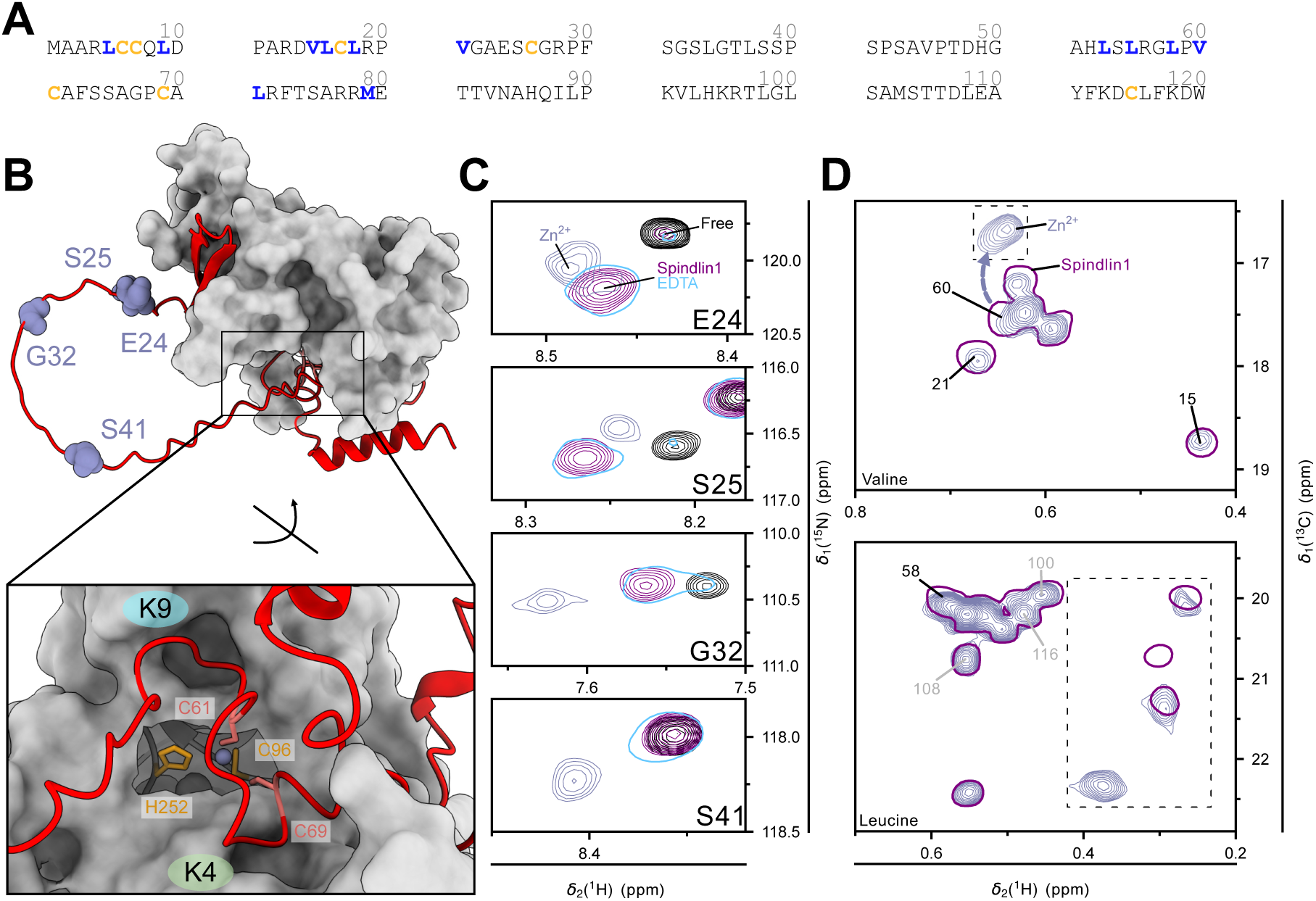
HBx_1-120_ and Spindlin1 form an inter-molecular zinc finger. **A.** HBx_1-120_ sequence with Cys residues in orange and methyl probes (Ile, Leu, Met, Val) in the two Spindlin1 interaction regions (residues 1–30 and 50–80) in blue. **B.** AF places a Zn^2+^ ion at the interface between HBx_1-120_ and Spindlin1 to form an inter-molecular zinc finger comprising C96 and H252 of Spindlin1 and C61 and C69 of HBx_1-120_; the latter two residues are suspected to form an intra-molecular zinc finger in full-length HBx_1-154_^17^. The interface on Spindlin1 is located between the two binding pockets that accommodate methylated K4 and K9 residues of histone H3 tails. AF predictions sample two alternative topologies of HBx_1-120_ (Fig. S4D). **C.** Excerpts from amide correlation spectra of free (black) and Spindlin1-bound HBx_1-120_ (purple; 1:1 ratio at 50 µM), after addition of Zn^2+^ ions (200 µM, grey), and supplemented with chelator EDTA (1 mM, blue). Residues from the linker region connecting the two interaction regions in HBx_1-120_ (Fig. 3D, E) display additional CSPs. The new CSP for S41, which is sandwiched between two Pro residues, could reflect a shift in cis/trans isomer distributions. All Zn^2+^-induced CSPs are fully reversible with EDTA; full spectra and intensity analysis in Fig. S4. **D.** Methyl correlation spectra of HBx_1-120_ bound to Spindlin1 before (purple single contour; 1:1 molar ratio at 50 µM) and after addition of Zn^2+^ ions (200 µM, grey). Dashed boxes are plotted at a lower counter level than the rest of the spectrum. Methyl probes near the zinc finger (C61 neighbours V60 and L58) disappear from the random-coil position. Four Leu signals tentatively assigned to the β-hairpin of HBx_1-120_ (Fig. 3E, F) display further CSPs upon Zn^2+^ addition; this effect is fully reversible upon addition of 1 mM EDTA (data not shown). Meanwhile, Leu residues outside the Spindlin1 binding region (grey labels) remain unchanged at the random coil position.

**Figure 5.**
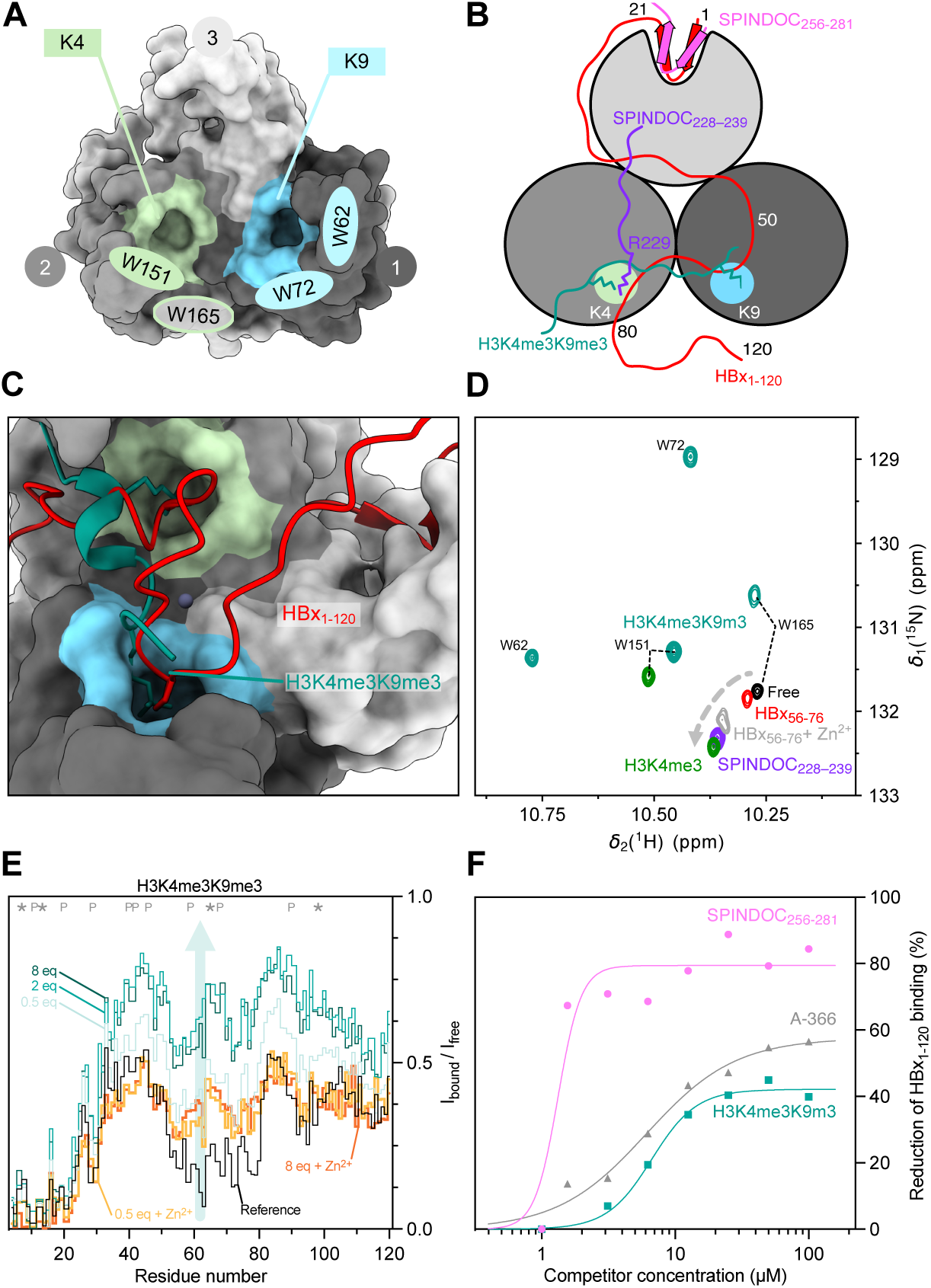
Methylated histone tails, SPINDOC and HBx_1–120_ compete for the same pockets on Spindlin1. **A.** AF model of Spindlin1 highlighting the pockets into which methylated or acetylated K4 and K9 residues of histone H3 tails insert. The K9 pocket is lined by W62 and W72; the K4 pocket is shaped by W151 and close to W165. **B.** Schematic of the competition for Spindlin1 between HBx_1-120_ (based on AF model), H3K4me3K9me3, SPINDOC_256-281_ and SPINDOC_228-239_ peptides (structures in Fig. S7). **C.** Superposition of AF-derived complex Spindlin1-HBx_1-120_-Zn^2+^ with the structure of H3K4me3K9me3 co-crystallized with Spindlin1 (PDB: 7CNA^40^) reveals direct competition for the same binding site. **D.** Excerpt from the Trp sidechain region of amide correlation spectra of Spindlin1 free and bound to peptides H3K4me3, H3K4me3K9me3, SPINDOC_228-239_ and HBx_56-76_ with/without Zn^2+^ (200 µM Spindlin1, 3 equivalents of peptide, 400 µM Zn^2+^). All peaks were assigned by mutagenesis. Only W165 is visible in free Spindlin1; W151 becomes visible upon binding H3K4me3. HBx_56-76_ and SPINDOC_228-239_ cause only small CSPs to W165 as they do not insert methylated Lys sidechains into the binding pockets; the CSP induced by HBx_56-76_ peptide approaches that of SPINDOC_228-239_ only in the presence of Zn^2+^ ions, indicative of a stabilization of the complex. Full spectra of backbone amide and His sidechain correlations are shown in Fig. S8 and S9. **E.** Amide intensity analysis shows that the histone-competing region (residues 50–80) of HBx_1-120_ is displaced from a complex with Spindlin1 (1:1 ratio at 50 µM) by two equivalents of H3K4me3K9me3 peptide, while leaving unaffected the N-terminal β-hairpin (residues 1–30), which binds the third Tudor barrel of Spindlin1. In the presence of Zn^2+^ ions, HBx_1-120_ competes more effectively against the histone peptide. In all competition experiments, intensity drops in the regions 50–60 and 70–80 are observed, at up to eight equivalents of H3K4me3K9me3 as well as in the presence of Zn^2+^. Analogous experiments with H3K4me3 peptide are shown in Fig. S6C. **F.** In the ELISA assay, the SPINDOC_256-281_ peptide causes a strong reduction in HBx_1-120_ binding to Spindlin1-coated plates, in line with NMR results (Fig. S6A). In contrast, H3K4me3K9me3 and A-366, a potent inhibitor of the Spindlin1-H3K4me3-interaction^47^, show weaker effects as they only compete against residues 50–80 of HBx_1-120_ for the secondary binding site on Spindlin1 (cf. panel E). The low HBx_1-120_ concentration in this assay (∼ 400 nM, 100 nM Spindlin1) and the use of milk powder as a blocking reagent imply the presence of Zn^2+^ ions in this setup. Fig. S10 details the validation of the ELISA assay.

### HBx_1-120_ and histone tails compete for the same site on Spindlin1

Having established that HBx_1-120_ engages Spindlin1 via an inter-molecular zinc finger, we explored whether this motif plays a functional role in transcriptional regulation. Spindlin1 recognizes methylated and acetylated histone tails^44^, epigenetic marks that regulate transcription by recruiting cofactors to host chromatin and, in the context of HBV infection, to the cccDNA^19^. Tri-methylation of K9 on histone 3 (H3K9me3) marks a repressive chromatin state, double tri-methylation at K4 and K9 (H3K4me3K9me3) signals a poised state, and tri-methylation at K4 alone (H3K4me3) promotes transcription^40,45^.

In our AF model, the inter-molecular zinc finger completed by HBx_1-120_ blocks access to the two Spindlin1 pockets that accommodate the methylated K4/K9 residues (Fig. 5B-C). Binding of HBx and histone tails to Spindlin1 would therefore be mutually exclusive. For experimental verification, we performed competition experiments with peptides covering the minimal recognition elements from histone tails (H3_1-20_ with K4me3 and K9me3 marks). In this NMR setup, recovery of signal intensity indicates that a peptide displaces the affected region of HBx_1-120_ from the Spindlin1 surface (Fig. 5E). Similar results were obtained with H3K4me3 and H3K4me3-K9me3 peptides (Fig. 5E and S6). In the absence of Zn^2+^, both peptides efficiently displace residues 50–80 of HBx_1-120_ from Spindlin1. Meanwhile, the N-terminal β-hairpin region displays non-competitive binding, which persists even after the histone-competing region has been displaced from the complex. Interestingly, the competition saturates with two equivalents of peptide while still exhibiting subtle intensity loss, indicative of residual transient interactions. In the presence of Zn^2+^, HBx_1-120_ competes effectively against up to an eightfold excess of peptide, indicative of a stronger HBx_1-120_-Spindlin1 interaction.

Complementary to the observations on HBx_1-120_, we performed Spindlin1-detected NMR experiments. The histone tail binding pockets of the first two Tudor domains are lined with aromatic side chains of His and Trp residues (Fig. 5A, Fig. S7D), which give rise to well-resolved NMR signals (Fig. 5D and S9A-B). The insertion of K4me3 or K9me3 side chains induces large CSPs. In comparison, the effects of binding the HBx_56-76_ peptide remain subtle, even in the presence of Zn^2+^, in line with the AF model in which HBx_1-120_ residues P59 and P68 occupy—but do not deeply insert into—the binding pockets (Fig. 5C). Overall, our NMR experiments prove that Zn^2+^ not only binds to the HBx_1-120_-Spindlin1 complex but also modulates its properties in a biologically relevant manner. The Zn^2+^ ion tightens the interface between Spindlin1 and HBx_1-120_, thereby more effectively shielding the functional surfaces on Spindlin1 against the very tight (low nM range^42^, Tab. S4) interaction with epigenetically marked histone tails. ITC (isothermal titration calorimetry) experiments, while providing quantitative affinities^24^, have so far been limited to short HBx-peptides, in part because full-length HBx is prone to aggregation. Our findings raise caution in interpreting relative affinities obtained for HBx fragments in isolation, which cannot fully capture the complexity of the interplay between various binding regions.

SPINDOC is a regulator of Spindlin1, for which a range of context-dependent effects on transcription were reported, from suppressive^46^ to selective activation of transcription^40^, specifically the differentiation between H3K4me3K9me3 and H3K4me3 marks^40,42^. Residues 1– 30 of HBx_1-120_ mimic the binding mode of SPINDOC on the third Tudor domain, completing the β-barrel (Fig. S7C). The effect of SPINDOC binding on NMR spectra of Spindlin1 was by investigated with two SPINDOC-derived peptides covering residues 228-239 or 256-281.

The SPINDOC_228-239_ peptide targets the histone-competing region of HBx_1-120_ (Fig. S7B). Like the HBx_56-76_ peptide, SPINDOC_228-239_ induces only subtle CSPs for Trp signals of Spindlin1 (Fig. 5D), consistent with a less immersive binding mode into the aromatic cages. NMR competition experiments confirm that HBx_1-120_ cannot compete against SPINDOC_256-281_ peptide for the third Tudor domain (Fig. S6), in line with previous ITC results^24,42^ (Tab. S4). Structurally, HBx_1-120_ mimics SPINDOC and may act as its antagonist or subvert its function. Quantitative assessment of the competition awaits evaluation with full-length SPINDOC protein.

Additionally, to enable a high-throughput screen for HBx_1-120_ interactors, we established an ELISA-based assay (Fig. S10). Competition for Spindlin1 between HBx_1-120_ and peptides or inhibitors can be reproduced using only a fraction of the protein quantity required for NMR experiments (Fig. 5F). However, a crucial advantage of the NMR-based assay is its ability to identify binding at the level of individual residues, so that competition can be detected even if only a subset of interaction sites is affected and overall complex formation is not abrogated.

## Discussion

We conducted an NMR study of conformational dynamics of the isoform HBx_1-120_, which originates from viral DNA fragments that integrate into the human genome in the context of chronic HBV infection^14^. We establish that HBx_1-120_ in isolation exists in a dynamically disordered state and undergoes folding upon binding human host proteins, here Bcl-xL and Spindlin1. Our results provide molecular-level evidence that HBx_1-120_ can bridge host proteins with non-overlapping binding regions to form ternary complexes, thereby enabling the hijacking of host cellular pathways, as previously demonstrated functionally for the DDB1-Smc5/6-HBx complex^18^.

Interactions of HBx_1-120_ can be monovalent and localized, as in the case of apoptosis regulator Bcl-xL, or multivalent, as observed with the epigenetic reader protein Spindlin1. The affinity for Bcl-xL is so low that this interaction is unlikely to be responsible for the carcinogenicity of the HBx_1-120_ isoform. In contrast, an extended interface is formed as HBx_1-120_ wraps tightly around the three Tudor domains of Spindlin1, occluding functionally relevant sites. This interface is stabilized by an inter-molecular zinc finger formed jointly by Spindlin1 and HBx_1-120_. Similar motifs, although rare, have been described before with proteins of eukaryotic origin involved in circadian^48^ and hormone^49^ regulation, immunity^50^ and oxidative stress response^51^ (cf. example in Fig. S5C). Extensive search of the Protein Data Bank (cf. Methods section) identified only a single occurrence of a virus-host protein complex jointly coordinating Zn^2+^: the regulatory Tat protein from human immunodeficiency virus 1 contributes three Cys residues to an inter-molecular zinc finger that targets Cyclin-T1 within the transcriptional elongation factor complex^52^.

The absence of a stable tertiary structure of HBx_1-120_ hinders high-resolution cryo-EM and has limited crystallization to short HBx fragments. Our strategy combines NMR with structure prediction to clarify where and how HBx targets host proteins. The multivalent binding between Spindlin1 and HBx_1-120_ highlights the importance of investigating integral HBx isoforms. Thanks to avidity, the full HBx_1-120_ isoform provides not only tighter binding but also achieves regulatory functions that have eluded prior investigations. For the first time, we succeeded in studying a large segment of HBx protein without tags or fusions in a setup capable of detecting even low affinity, dynamic complexes.

HBV is a highly evolved and minimalistic virus^3^. The HBx protein compresses multiple interaction motifs into a short primary structure. In addition to the viral strategy of mimicking linear interaction motifs from host proteins (in HBx: β-hairpin of SPINDOC, histone tails, BH3-like helices from Bcl family, H-box of DDB1 and CUL4-associated factors), we here discovered an inter-molecular zinc-binding motif. This crucial element targets directly the histone docking region in Spindlin1 with just two key amino acid residues as a ‘sticky patch’. Given the extraordinary abundance of Cys residues in HBx (Fig. 6B), it is conceivable that inter-molecular zinc-binding motifs may drive other host protein interactions as well. This motif requires strategically placed His or Cys residues on Spindlin1, hinting at the existence of a human protein that uses the same binding mode. Cys residues could provide further gain of function through dimerization^53^ or intra-molecular zinc fingers (Fig. 6C).

**Figure 6.**
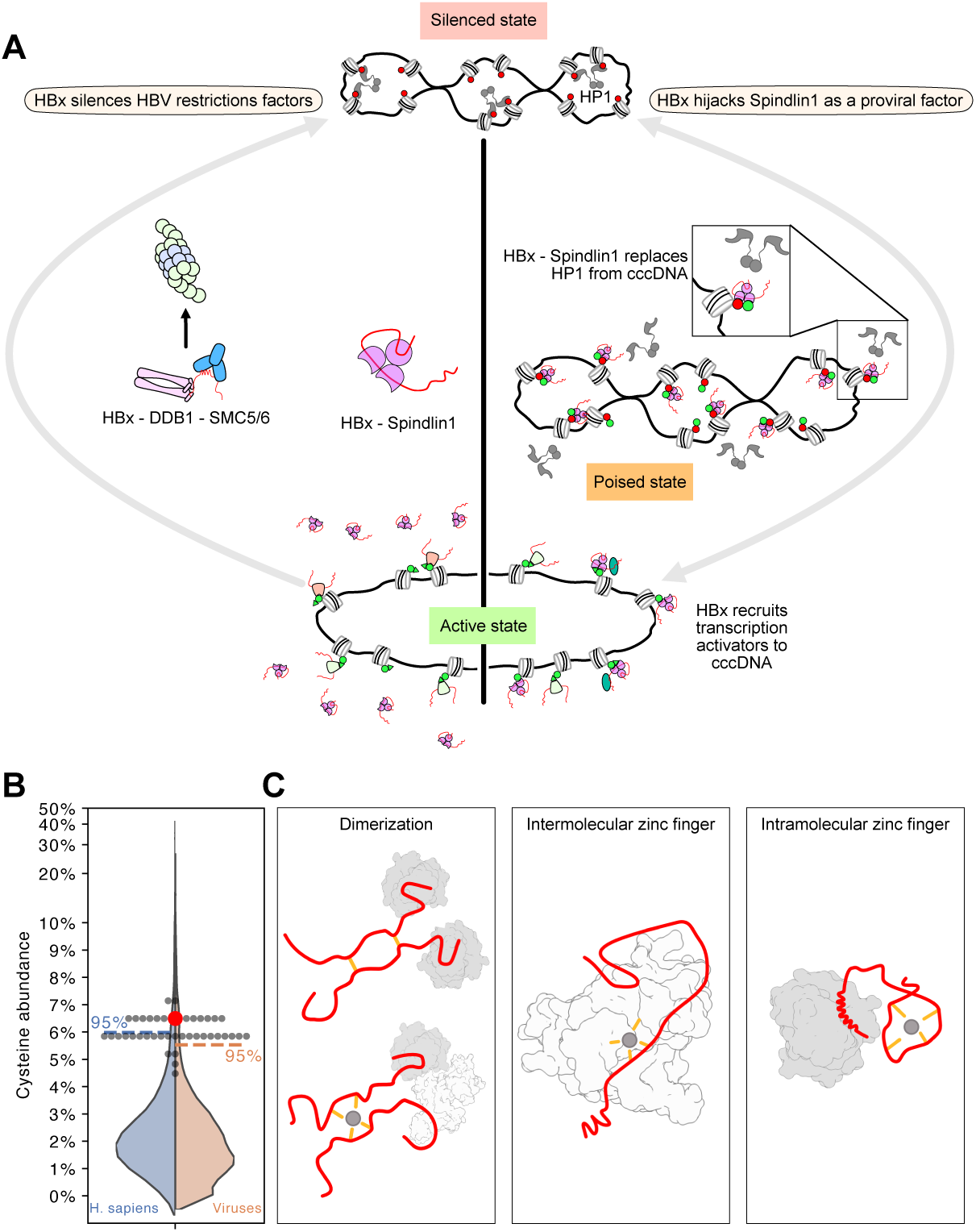
HBx counteracts silencing of the cccDNA. **A.** Two mechanisms were deduced from in-cell experiments, both ultimately result in active cccDNA decorated with H3K4me3 and hyperacetylation marks. *Left*: Ducroux et al.^19^ report an increase in silencing H3K9me3 marks and Spindlin1 recruitment to the cccDNA in the absence of HBx. HBx masking Spindlin1 (pink complex establish by NMR in this work) blocks it from the cccDNA. This mechanism resembles how HBx degrades the restriction factor SMC5/6 by hijacking of the DDB1 ubiquitin ligase complex. Despite the observation of a ternary complex HBx-Spindlin1-DDB1 *in vitro*^25^, Ducroux et al. did not observe variations in overall Spindlin1 levels in the presence of HBx, suggesting no involvement of proteasomal degradation pathways. In sum, this model proposes that HBx blocks every restriction that otherwise would silence the cccDNA. *Right*: Liu et al. propose a model analogous to the interaction between Spindlin1 and its regulator SPINDOC. As the cccDNA enters the cell, host defence mechanisms decorate it with H3K9me3 marks, prompting HP1 dimers to compact the cccDNA. To reactivate the cccDNA, HBx mimics SPINDOC and increases the affinity of Spindlin1 for bivalent H3K4me3K9me3 marks. Indeed, the Spindlin1-SPINDOC complex can remove HP1 proteins through competition^40^. The transition from silenced to poised state requires the action of another player, which could be recruited by HBx. In sum, HBx liberates the silenced cccDNA and recruits further transcription activators. This model assumes a dynamic equilibrium between the silenced and active state. **B.** Statistics of Cys residue abundance in proteins belonging to the virus or *Homo sapiens* taxonomy. HBx genotype D sequence from our experiments is indicated in red, other human and ape genotypes are indicated in black. All HBx sequences are ranked in the top 7% or 5% of viral and human proteins’ Cys abundance, respectively. **C.** Illustration of the different mechanisms by which Cys residues can enable short IDPs to form diverse interactions. Dimerization of HBx through disulfide bonds or zinc fingers has been reported before without any link yet to a specific pathway or interaction^26,53^. This study identifies an inter-molecular zinc finger that stabilizes the interface between the HBx_1-120_ isoform and Spindlin1. Full-length HBx_1-154_ is suspected to form an intra-molecular zinc finger^17^, which conceivably could stabilize the HBx segments not directly involved in the interaction, as observed in epigenetic regulators of human origin^62^.

HBx is considered a driver of liver disease and HCC^13,14^. It interferes with epigenetic regulation^54^ to promote the expression of viral^55^ and host genes^56^. The interaction between HBx_1-120_ and Spindlin1 subverts cellular mechanisms that otherwise would counteract transcription from the extrachromosomal cccDNA^54^. On the molecular level, we establish that HBx_1-120_ occludes docking sites on Spindlin1 that recognize epigenetically marked histone tails; HBx further competes with the Spindlin1 regulator SPINDOC (C11orf84) through molecular mimicry.

Previously, two cell biological studies^19,24^ proposed partially conflicting mechanisms of the role of Spindlin1 in HBV infection (Fig. 6). As final consequence of HBx expression, both studies report the increase of H3K4me3 and acetylation marks at the cccDNA, which signals a transcriptionally active state to benefit viral replication. Yet the proposed mechanism by which HBx achieves this differ: According to Ducroux et al.^19^, based on an in-cell ChIP experiment post infection in hepatocyte culture, Spindlin1 emerges as a restriction factor of HBV. Spindlin1 depletion is correlated with an increase of active marks on cccDNA. The proposed mechanism is reminiscent of HBx hijacking DDB1 to target the HBV restriction factor Smc5/6 for degradation, except that with Spindlin1, HBx merely masks and does not deplete the restriction factor. In the presence of HBx, Spindlin1 is not recruited to the cccDNA (Fig. 6A). A similar regulatory mechanism targeting histone interactions has been reported for human proteins^57^. Ducroux et al. also observe a similar effect with other DNA viruses such as HSV-1. While it may appear counterintuitive that HBx would silence the putative transcriptional activator Spindlin1, in this scenario, increased H3K4me3 and acetylation marks on cccDNA would be accomplished by another epigenetic regulator.

On the contrary, Liu et al.^24^ report that HBx hijacks Spindlin1 to promote transcription. In a similar experimental setup to the previous, the authors observe increased recruitment of Spindlin1 to the cccDNA under overexpression of HBx. Once in the cell, the cccDNA is epigenetically silenced by an ensemble of regulators, such as SETDB1 depositing H3K9me3 marks, which signal chromatin compactors (HP1 heterochromatin protein) to block transcription. A poised state is invoked with cccDNA decorated with bivalent H3K4me3K9me3 marks. In ITC experiments, Liu et al. observe that a complex between Spindlin1 and HBx_2-21_ peptide has higher affinity for bivalent compared to monovalent K4me3 marks. A similar model was proposed for SPINDOC regulating Spindlin1^40^, in line with the notion that HBx mimics SPINDOC. In light of the second histone-competing region in HBx identified by NMR, affinities would need to be revisited with larger segments of HBx. By redirecting Spindlin1 towards chromatin with bivalent marks, HBx would contribute to the removal of repressors such as HP1 and to the recruitment of activators, ultimately achieving an active state of the cccDNA (Fig. 6A).

The two studies observe increased ^24^ versus decreased ^19^ recruitment of Spindlin1 to the cccDNA in the presence of HBx. This discrepancy may be linked to differences in experimental setup, notably the use of HepaRG *vs*. HepG2 cells, which differ in gene expression^58^. The discrepancy also reflects the complexity of observing switch-like epigenetic regulation over fluctuating viral infection cycles. Our structural model is in line with the mechanism proposed by Ducroux et al.: The zinc finger on the surface of Spindlin1 is positioned right between the pockets for H3K4me3 and H3K4me3K9me3; HBx_1-120_ competes against both. Nevertheless, the observation that HBx can be easily outcompeted by the SPINDOC_256-281_ hairpin questions its capacity to fully restrict Spindlin1. Moreover, HBx would need to incapacitate multiple other epigenetic regulators. Our structural model is also connected to the hypothesis by Liu et al.: the intermolecular zinc finger increases the energy barrier for binding histone tails, redirecting the HBx-Spindlin1 complex towards more tightly binding bivalent marks and thus favouring poised cccDNA. We could not observe a significant difference in the ability of H3K4me3 and H3K4me3K9me3 peptide to replace HBx_1-120_ from Spindlin1 in our NMR setup, though. In summary, we clearly establish that HBx targets Spindlin1 to disrupt epigenetic signalling.

When Spindlin1 activates oncogenic transcriptional programs, even outside the context of HBV infection, it can promote cell cancer proliferation, notably through the Wnt signalling pathway ^59,60^ and also under involvement of SPINDOC^61^. Improper distribution of Spindlin1 within chromatin caused by HBx could unleash transcriptional activators and activate silenced oncogenes in HCC patients. Unlike HBx_1-120_, full-length HBx_1-154_ can form an intra-molecular zinc finger according to AF (involving C61, 69, 115 and H137, Fig. 1B) and more tightly engage host proteins like Bcl-xL via the C-terminal 34 residues. Conversely, HBx_1-154_ might have a lower propensity for inter-molecular Zn^2+^ binding to hijack Spindlin1. Furthermore, the isoform of HBx could define which third proteins can be recruited into a ternary complex with Spindlin1, thereby rewiring the transcriptional landscape.

Achieving HBV cure by fully eliminating both the cccDNA and HBV DNA integrates from host chromatin poses a high bar. An easier approach could be to silence the cccDNA^10^. In this context, a molecular understanding of how HBx engages and subverts Spindlin1 is an important foundation to develop HBx as a therapeutic target.

## Supplementary Figures

**Figure S1.**
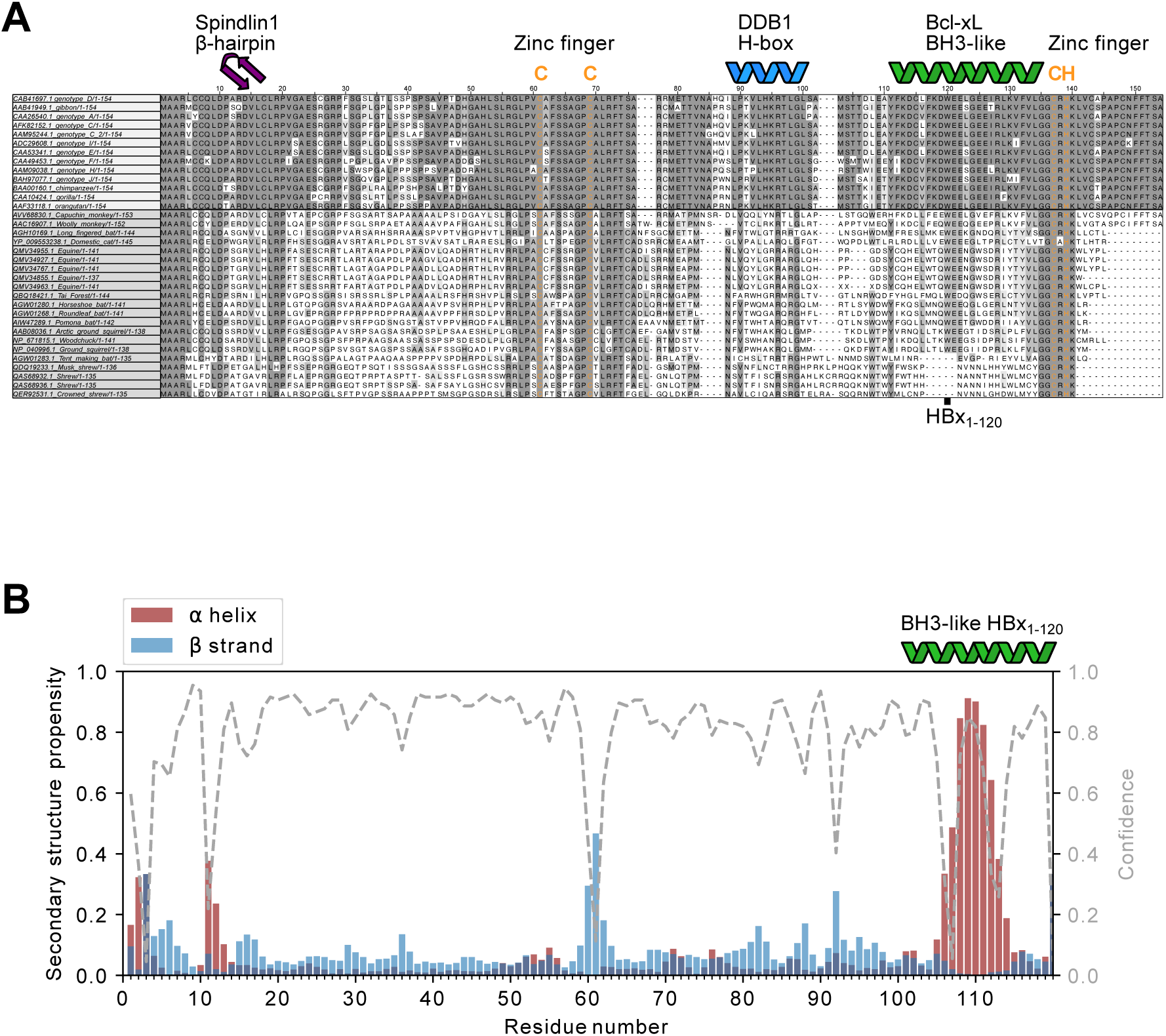
Sequence conservation of HBx and secondary structure propensity of HBx_1-120_. **A.** Multiple sequence alignment of HBx protein. Sequences are sorted into two groups: human genotypes and ape viruses (top) and other mammals (bottom) based on conservation of the sequences. Amino-acid residues are coloured based on conservation in their respective groups. The alignment shows the high conservation of the zinc finger CCCH motif and their respective neighbours, despite the sequences showing a diversity of lengths. Among human and ape HBx sequences, the least conserved region spans residues 30–40, which is predicted as a loop linking two binding regions in the HBx–Spindlin1 complex. **B.** Secondary structure predictions based on NMR chemical shifts obtained from TALOS-N^63^. The ^13^C chemical shifts of HBx_1-120_ could only be obtained in a buffer that prevents the protein from aggregating (1 M urea, 150 mM NaCl, 50 mM HEPES, 125 mM L-arginine, 5 mM DTT, pH 7.4). Despite the high urea content, the HBx_1-120_ protein shows a propensity to form a helix (BH3-like domain), which mediates interactions with Bcl family members^31^.

**Figure S2.**
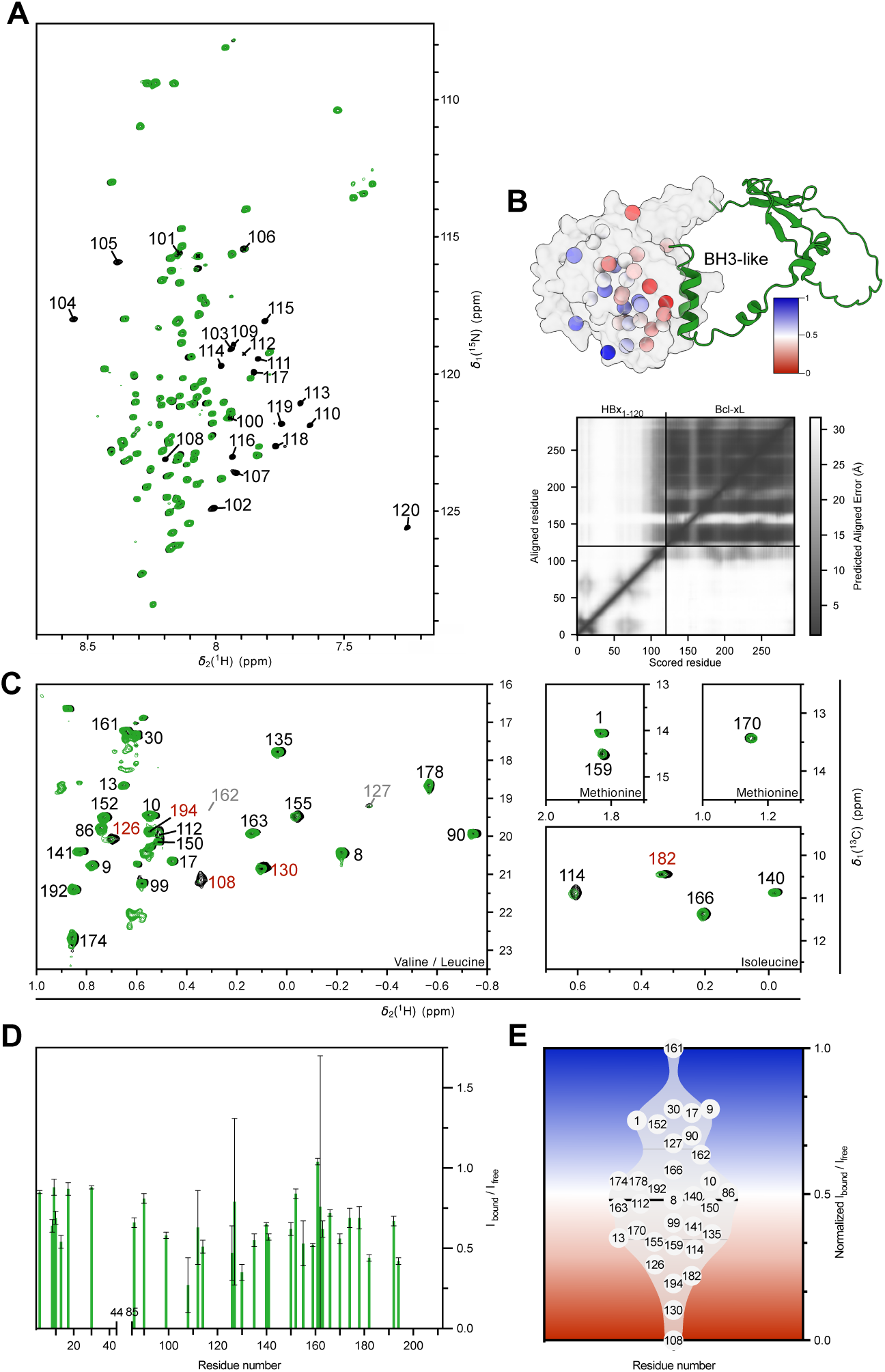
The interaction between HBx_1-120_ and Bcl-xL is monovalent. A. Amide correlation spectra (SOFAST ^1^H-^15^N HMQC) of HBx_1-120_ in free form (black) and bound to Bcl-xL (green; 50 µM, 1:1 ratio). Signals from residues 100–120 display complete intensity loss in the complex. All other signals accurately overlap with the reference, indicating that only a single domain folds upon binding Bcl-xL, while the remainder of HBx_1-120_ remains unfolded **B.** AF model of the complex between the Bcl-xL (surface representation; residues 1–44//85–212, cf. Methods section) and HBx_1-120_ (cartoon) with red indicating the strongest loss of methyl peak intensity (panel D) in the complex. The accompanying AF PAE matrix shows high confidence for the prediction of the C-terminal helix of HBx_1-120_ and its position relative to the BH3 groove of Bcl-xL. Confidence for the remaining fold of HBx_1-120_ and its orientation relative to Bcl-xL is low, consistent with the disordered state of HBx_1-120_ observed by NMR. **C.** Methyl correlation spectrum (^1^H-^13^C HMQC) of Bcl-xL (ILVM-^13^CH_3_-labelling) in its free form (black) and bound to HBx_1-120_ (green; 50 µM, 1:1 ratio). Assignments were transferred from reference^64^ with residue numbers according to the non-truncated Bcl-xL protein. Red labels indicate signals showing the highest intensity loss in the bound complex. **D.** Loss of methyl peak intensity from panel **C** (Met, Leu, Val, Ile) between free Bcl-xL and in complex with HBx_1-120_, expressed as the intensity ratio I_bound_/I_free_ **E.** Normalized intensity ratios (scaled to the [0–1] range) from panel D represented as a distribution plot. The background gradient indicates how residues are coloured in the structure in panel B.

**Figure S3.**
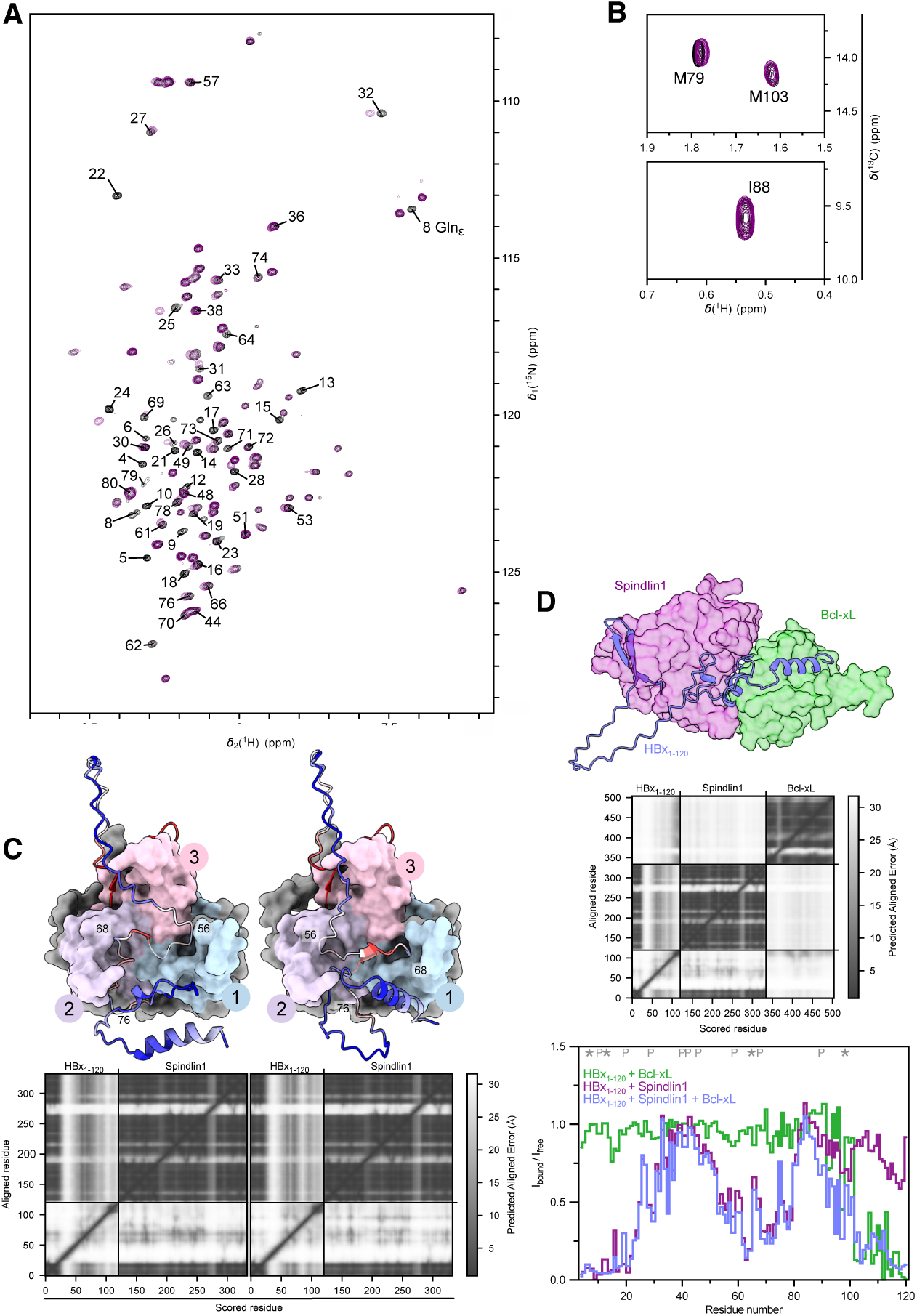
The interaction between HBx_1-120_ and Spindlin1 is bivalent. **A.** Amide correlation spectrum (SOFAST ^1^H-^15^N HMQC) of HBx_1-120_ in free form (black) and bound to Spindlin1 (purple; 50 µM, 1:1 ratio). Signals from residues in region 1–20, including the sidechain of Q8, show complete intensity loss in the complex, consistent with formation of a β-hairpin docking onto Tudor domain 3. Residues 20–30 exhibit CSPs, while residues 50–80 display partial signal attenuation, indicating a second binding region. **B.** Excerpts from methyl correlation spectrum (^1^H-^13^C HMQC) of HBx_1-120_ (ILVM-^13^CH_3_-labelling) in its free form (black) and bound to Spindlin1 (purple; 50 µM, 1:1 ratio). Complementary to data in Fig. 3F, the signal of M79 displays a slight shift, highlighting the second binding region of HBx_50-80_ on Tudor domains 1 and 2 of Spindlin1. Meanwhile, residues I88 and M103 show no perturbation. The signals of M79 and M103 were assigned by deduction, based on the spectral changes observed for Spindlin1 and Bcl-xL. **C.** AF predicts two alternative topologies for residues 50–80 on the surface of Spindlin1. The key difference lies in the order in which the histone-binding pockets are engaged. The N-terminal region of HBx_1-120_ consistently contacts Tudor 3. In the 3-1-2 model (left), HBx first docks onto Tudor 1 domain (K9me3 pocket) and then Tudor 2 (K4me3); reversed in the alternative 3-2-1 model (right). Experimental data satisfy both models, since the same residues contact the Spindlin1 surface in each conformation. The PAE matrix shows high confidence for the HBx_1-20_ hairpin region and low confidence for residues 50–80 covering the histone-binding region. **D.** Intensity analysis of amide correlations of HBx_1-120_, expressed as the intensity ratio I_bound_ /I_free_, was obtained from SOFAST ^1^H-^15^N HMQC spectra of the protein in free form, bound to Spindlin1, Bcl-xL alone, or both simultaneously (all at 50 µM; 1:1:1 ratio). Both binding partners cause additive intensity losses, consistent with the formation of a ternary complex at least transiently. AF also predicts a ternary complex. The PAE matrix shows high confidence for the positions of the dedicated binding partners but low confidence for Spindlin1 relative to Bcl-xL, supporting the notion that HBx bridges two proteins that do not interact in its absence. The interaction between Bcl-xL and Spindlin1 is unlikely to be biologically relevant, as the two do not colocalize in the cell.

**Figure S4.**
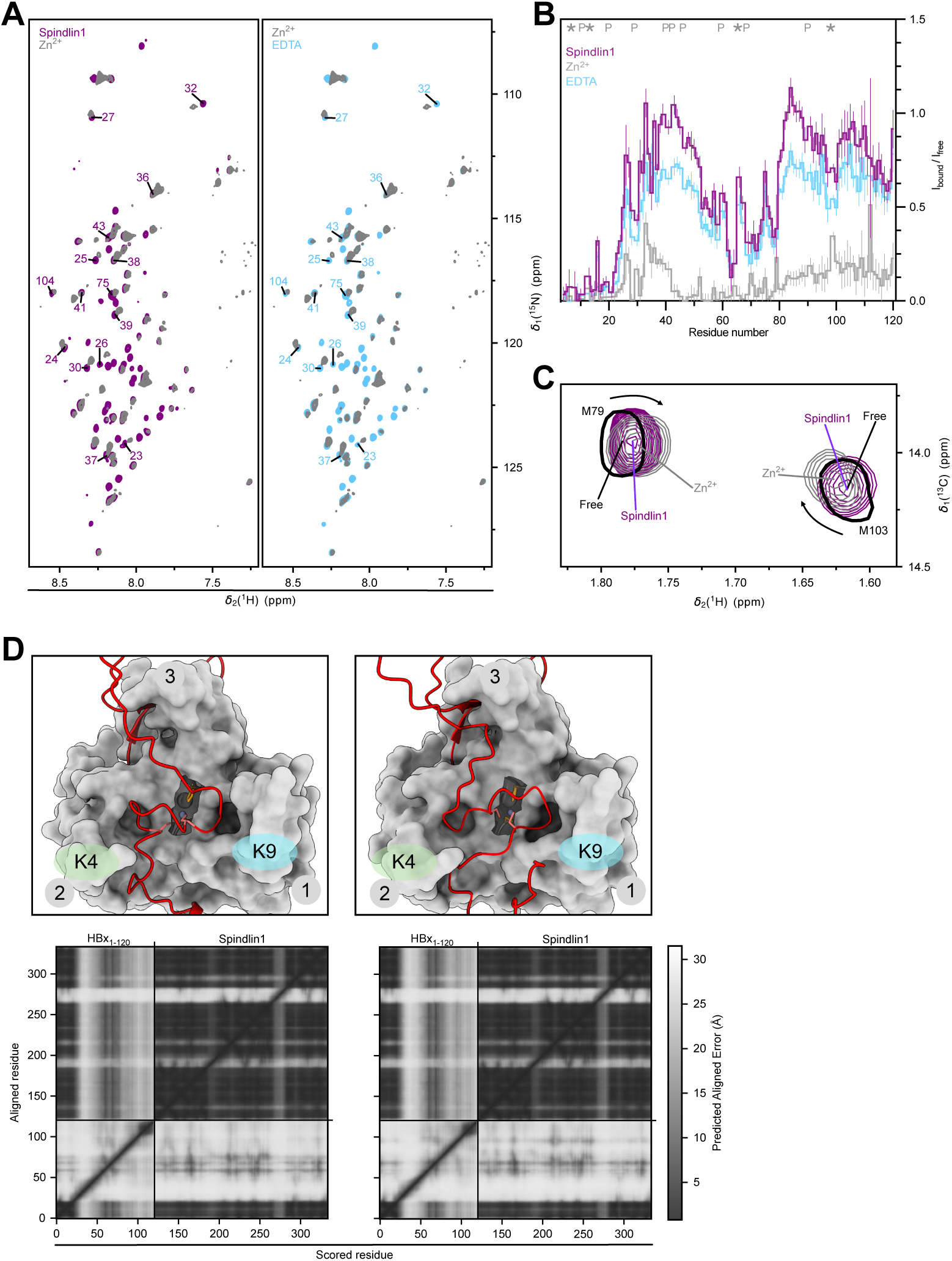
HBx_1-120_ and Spindlin1 form an inter-molecular zinc finger. **A.** Amide correlation spectra (SOFAST ^1^H-^15^N HMQC) of HBx_1-120_ in complex with Spindlin1 (purple; 50 µM, 1:1 ratio), supplemented with Zn^2+^ (400 µM, grey contours) and after addition of EDTA (blue, 1 mM). Zn^2+^ induces non-uniform intensity changes to the spectrum in addition to some CSPs (corresponding residues labelled), particularly to the loop connecting the two binding regions. The effect from Zn^2+^ addition is fully reversible by EDTA. **B.** Intensity analysis of amide correlations of HBx_1-120_ for spectra from panel A, expressed as intensity ratio I_bound_ / I_free_. A major intensity loss is observed upon Zn^2+^ addition, with residual signals primarily from residues outside the Spindlin1-interacting region. The slight loss of intensity between spectra recorded without Zn^2+^ and in the presence of Zn^2+^ and EDTA are attributable to minor protein loss during handling and dialysis, as well as gradual aggregation of HBx_1-120_ over time. **C.** Excerpts from methyl correlation spectra (^1^H-^13^C HMQC) of free HBx_1-120_ (black; ILVM-^13^CH_3_-labelling), bound to Spindlin1 (purple; 50 µM, 1:1 ratio) and in the presence of Zn^2+^ (grey; 400 µM). M79 shows a different CSP in the presence of Zn^2+^ compared to Spindlin1 alone; the CSP is reversed by EDTA (data not shown). **D.** Irrespective of the absence (cf. Fig. S3C) or presence of a Zn^2+^ ion, AF predicts two topologies of HBx_1-120_ that differ in the order in which it contacts the Tudor domains: 3-1-2 (left) and 3-2-1 (right). Residue 91–120 are omitted in both models to simplify structural representation.

**Figure S5.**
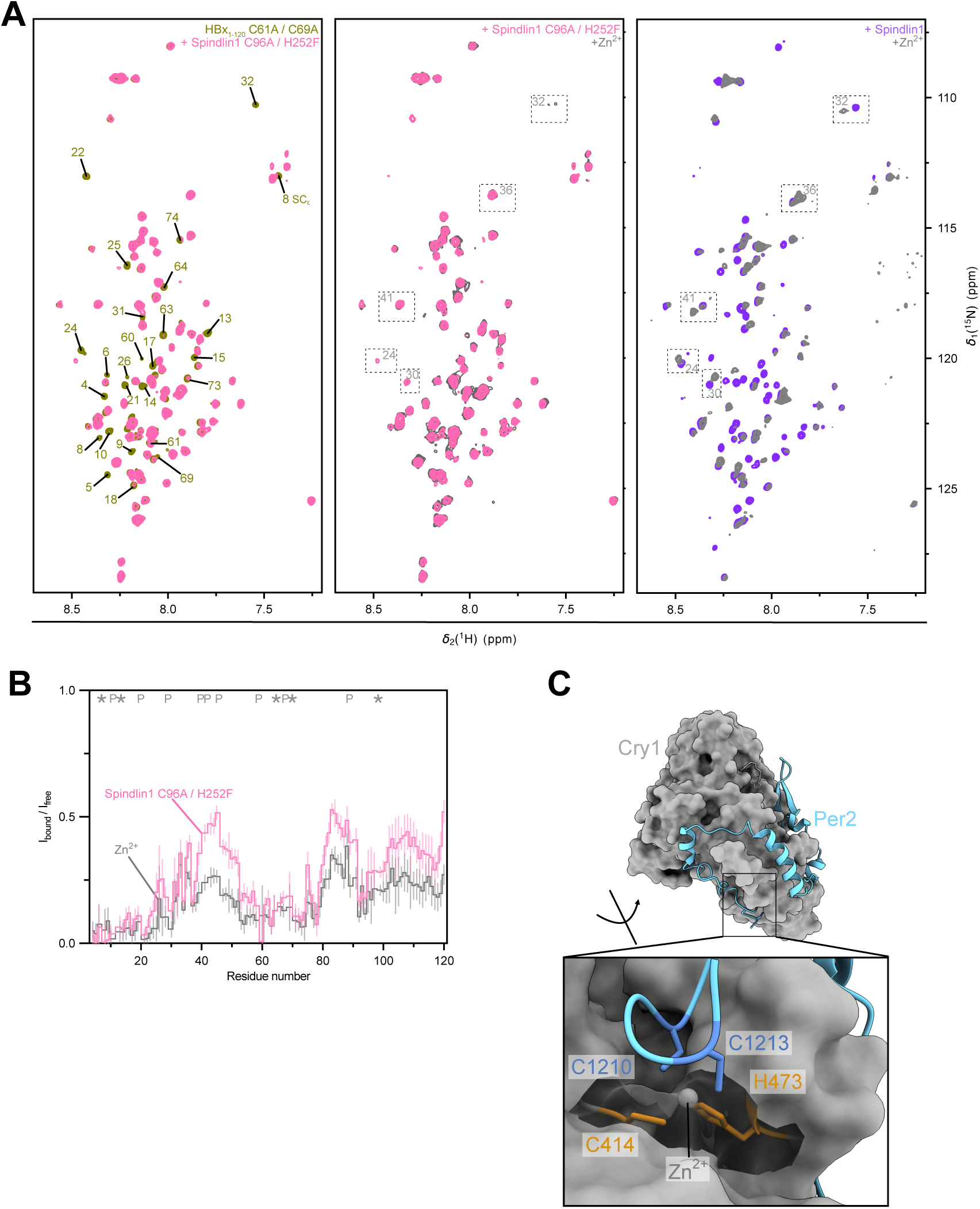
Removing the CCCH residues in HBx_1-120_ and Spindlin1 abolishes zinc-dependent binding. **A.** Amide correlation spectra (SOFAST ^1^H-^15^N HMQC) recorded on double mutant HBx_1-120_^C61A/C69A^ free and in complex with mutant Spindlin1^C96A/H252F^ (purple; 50 µM, 1:1 ratio) and supplemented with Zn^2+^ (200 𝜇M). Mutated residues were tentatively assigned. Mutation of all four zinc finger residues from CCCH to AAAF does not impair the binding of the partners in either binding region but abrogates its sensitivity to zinc. For comparison, analogous spectra of the wild-type protein highlight Zn-induced CSPs (grey boxes, same spectra as Fig. S4A). **B.** Intensity analysis of amide correlations of HBx_1-120_ obtained from spectra in panel A, expressed as intensity ratio I_bound_ / I_free_. A uniform intensity loss is observed after addition of Zn^2+^, which is caused by slight reduction of protein concentration during Zn^2+^ dialysis, sample handling and gradual aggregation of HBx_1-120_ C61A/C69A over time. The uniform intensity loss contrasts the observation with the wild-type proteins (Fig. S4B), where a pronounced intensity loss is specific to the two binding regions while signals from the linker are less attenuated and display CSPs instead. **C.** The interface between circadian clock proteins CRY1 (cryptochrome circadian regulator 1) and PER2 (period circadian protein homolog 2) is mediated by an inter-molecular CCCH zinc finger. Mutation of residues involved modulates the strength of the interaction.^48^

**Figure S6.**
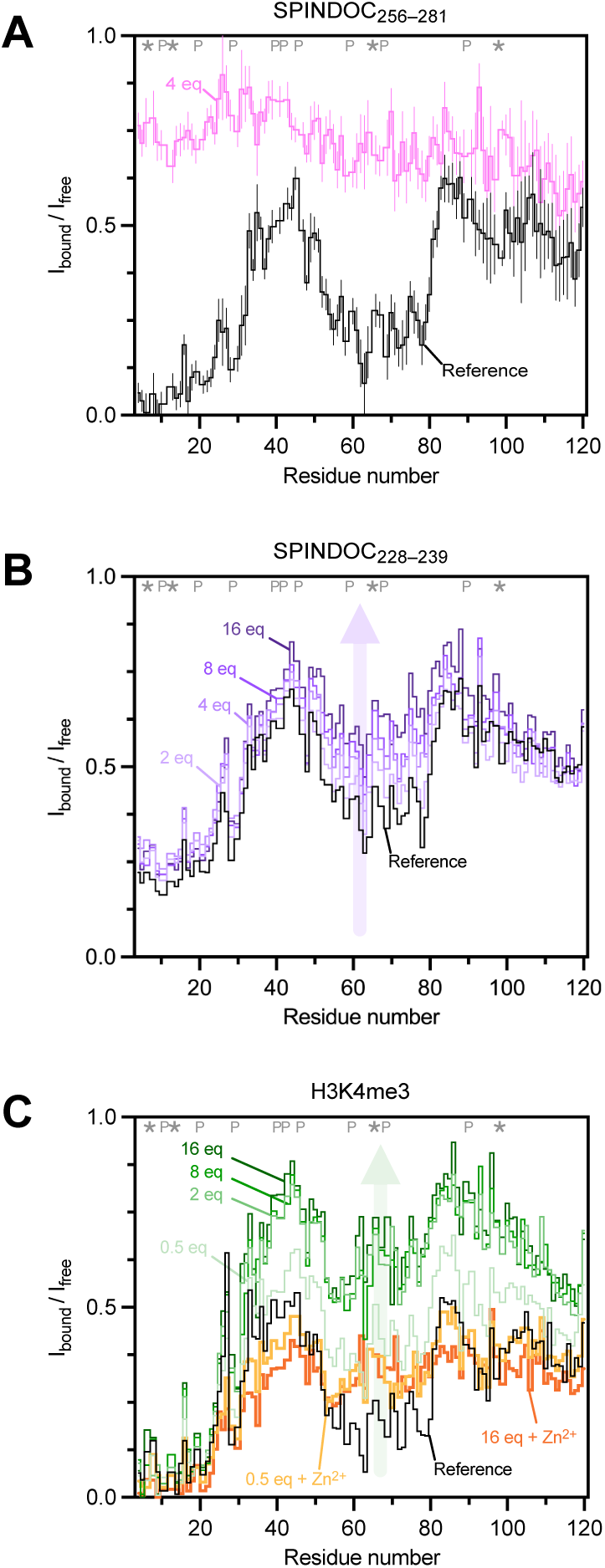
Competition for Spindlin1 between HBx_1-120_ and peptides derived from SPINDOC and histone tails. The intensities of amide correlations, I_bound_ / I_free_, were obtained from SOFAST ^1^H-^15^N HMQC spectra of HBx_1-120_ in complex with Spindlin1 (black reference) and after addition of the respective peptides (coloured, 1 eq equals 50 µM). Competing regions are visualized structurally in Figs. 5E and S7. **A.** Competition assay with SPINDOC_256-281_. This peptide docks as a hairpin onto the Tudor 3 domain of Spindlin1 and fully outcompetes HBx_1-120_ at four equivalents. Binding to the second, non-competitive HBx_1-120_ region is also abrogated, confirming that the multivalent interaction is driven by the β-hairpin. **B.** Competition assay with SPINDOC_228-239_. This peptide competes for the Tudor 2 domain on Spindlin1 where it occupies the K4me3 binding pocket with an Arg sidechain. According to the AF model (Fig. S7B), the competition is mild as SPINDOC_228-239_ does not directly occupy the same area as HBx_1-120_. Even 16 eq of peptide do not fully outcompete the second interaction region of HBx_1-120_. The slightly higher base level of the reference compared to experiments in panels A and C is due to imperfect complex formation or stoichiometry between HBx_1-120_ and Spindlin1 even before the peptide titration. **C.** Competition assay with H3K4me3 peptide (residues 1–20 of histone H3). This peptide competes for Tudor domain 2 of Spindlin1. Upon addition of 2 eq of peptide, HBx_1-120_ is mostly displaced from this binding region; subtle local intensity losses remain around residues 50–60 and 70–80. Stabilizing the Spindlin1-HBx_1-120_ complex with 200 µM Zn^2+^ resulted in more effective competition; 0.5 and 1 eq of peptide resulted in similar intensity loss compared to the reference.

**Figure S7.**
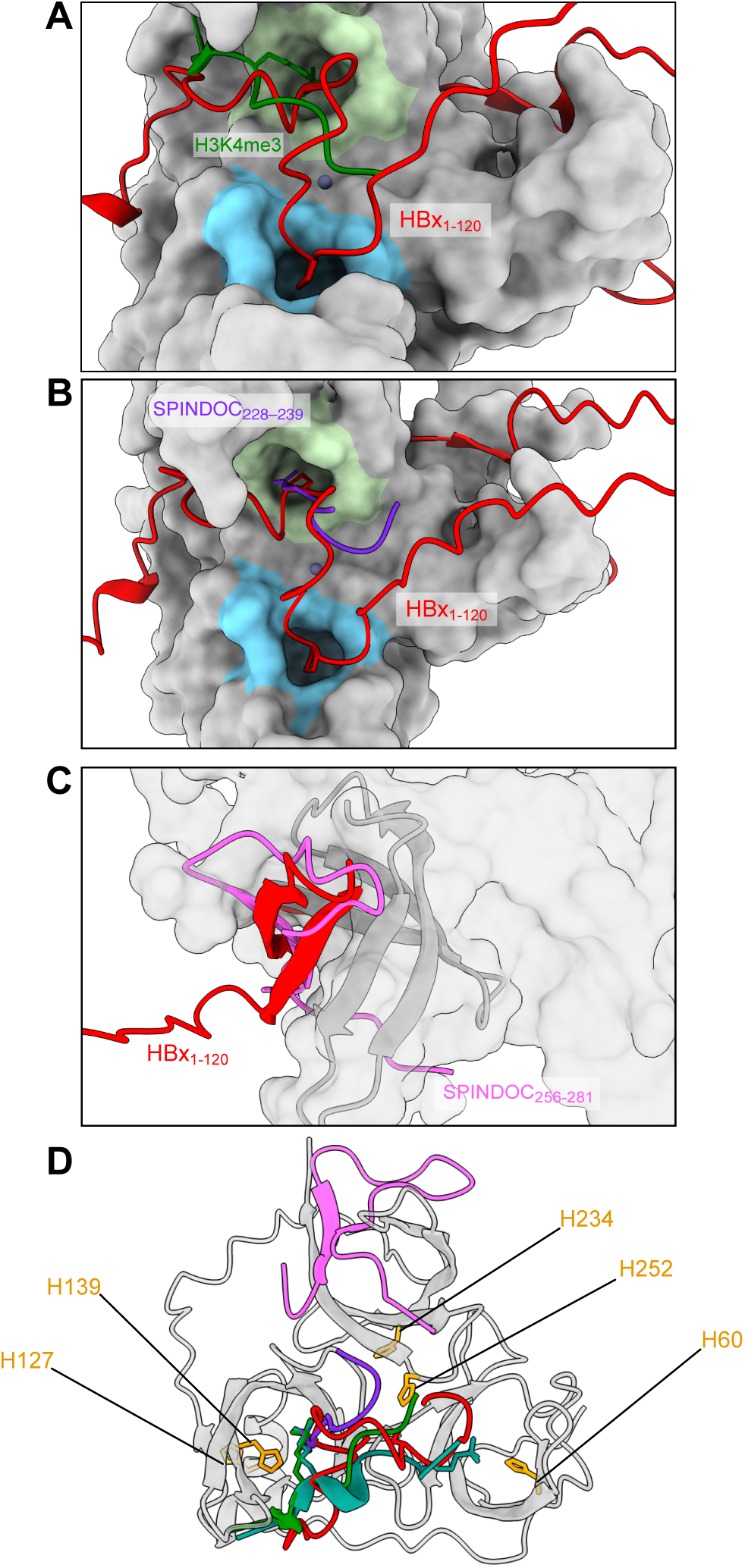
HBx_1-120_, SPINDOC and histone tails compete for the same binding pockets on Spindlin1. **A.** AF model of HBx_1-120_-Spindlin1 complex in the presence of one Zn^2+^ ion superimposed with the crystal structure of H3K4me3 histone-derived peptide bound to Spindlin1 (PDB 4MZG^65^). Similarly to H3K4me3K9me3 (Fig. 5C), this peptide competes against residues 50–80 of HBx_1-120_ for the same binding site. K4me3 and K9me3 binding pockets are indicated in green and blue, respectively **B.** AF model of HBx_1-120_-Spindlin1 complex in the presence of one Zn^2+^ ion superimposed with the crystal structure of SPINDOC_228-239_ (Docpep2) bound to Spindlin1 (PDB 7EA1^42^). While the peptide is not fully resolved in the crystal structure; it clearly competes partially against HBx_1-120_ for the histone-tail binding region. The peptide inserts the side chain of R239 into the K4me3 binding pocket; in the AF model HBx_1-120_ docks to the same binding pocket with P58 and P69 residue. **C.** AF model of HBx_1-120_-Spindlin1 complex in the presence of one Zn^2+^ ion superimposed with the crystal structure of SPINDOC_256-281_ (Docpep3) (PDB 7E9M^42^). SPINDOC_256-281_ competes directly against the N-terminal region of HBx_1-120_ for the Tudor 3 domain of Spindlin1. HBx and SPINDOC adopt a similar β-hairpin fold to complete the Tudor β-barrel. **D.** Distribution of His residues in Spindlin1 is visualized on an AF model of HBx_1-120_-Spindlin1 complex together with the crystal structures of peptides SPINDOC_228-239_, SPINDOC_256-281_, H3K4me3, and H3K4me3K9me3. H252 mediates the intermolecular zinc finger; the side chains of H60 and H139 act as NMR probes of binding partner entering the K4me3 and K9me3 pockets.

**Figure S8.**
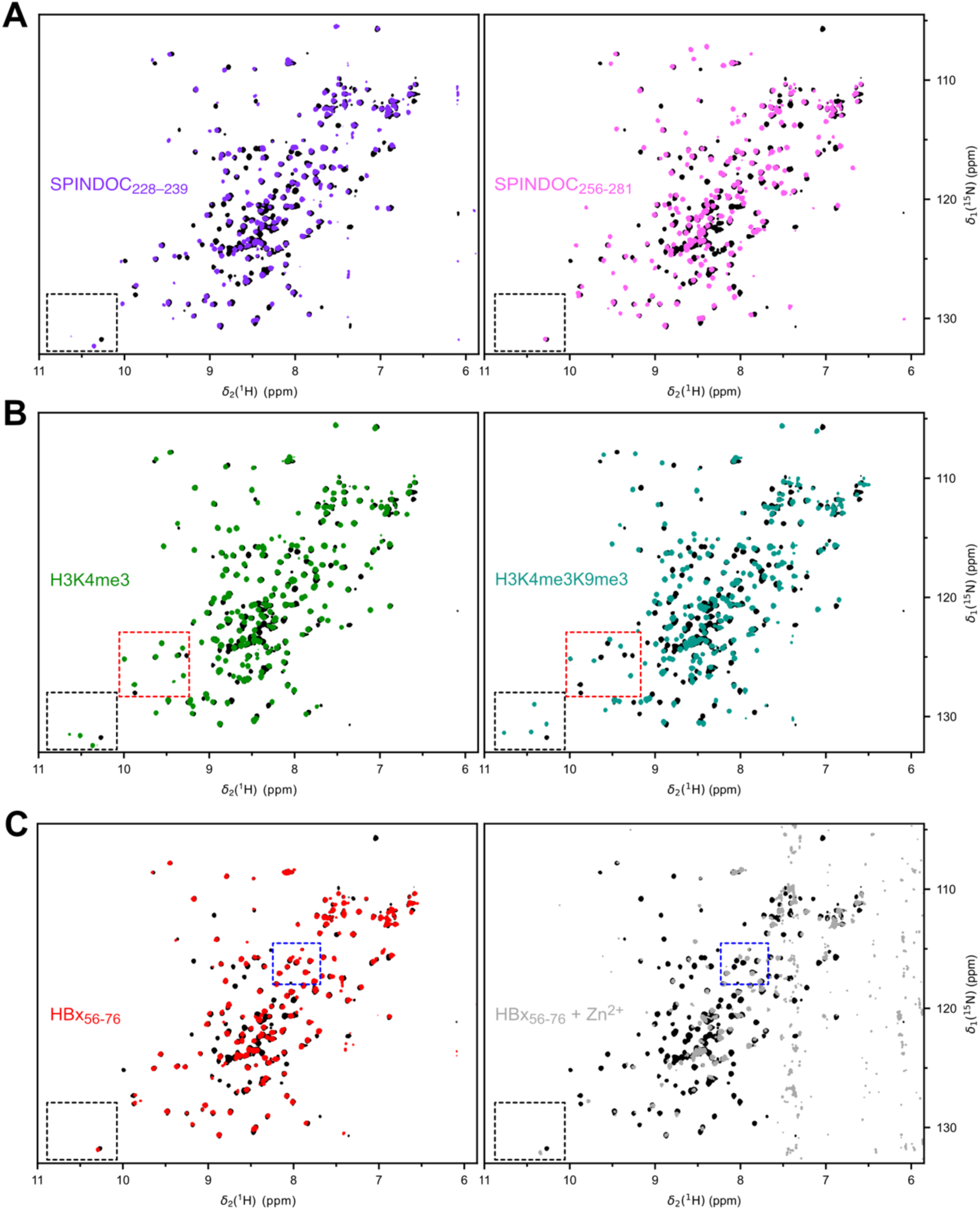
Spindlin1-detected NMR binding experiments of peptides from HBx, SPINDOC and histone tails. Amide correlation spectra (_1_H-15N SOFAST HMQC) of Spindlin1 free (black) and bound to the respective peptides from histone tails, SPINDOC and HBx (coloured). A. Spectra in the presence of SPINDOC_228-239_ and SPINDOC_256-281_ (200 µM Spindlin1, 1:3 ratio). The two peptides interact with different Tudor domains, which reflects in the spectra as intensity losses and CSPs for different sets of signals. Overall, SPINDOC_256-281_ has a larger impact, as the docking of the β-hairpin onto Tudor 3 domain causes a significant conformational change to Spindlin1. SPINDOC_228-239_ displays a pronounced CSP on the W165 sidechain (box) as its arginine sidechain enters the K4me3 pocket. B. Spectra in the presence of histone-derived peptides H3K4me3 and H3K4me3K9me3 (200 µM Spindlin1, 1:3 ratio). Both peptides target similar surfaces on Spindlin1; however, H3K4me3K9me3 has a larger impact (cf. red box), as it spans both Tudor domains and can rigidify Spindlin1 by bridging two binding pockets. Both peptides trigger CSP for the Trp sidechains (black box) that line the respective binding pockets (Fig. 5A). C. Spectra in the presence of HBx_56-76_ with and without Zn^2+^ (200 µM Spindlin1, 1:3 ratio, 400 𝜇M Zn^2+^). HBx_56-76_ peptide induces intensity loss of a subset of signals distinct from other studied peptides. The addition of Zn^2+^ increases the impact on the spectrum, with more defined intensity loss and CSPs (blue boxes). This effect is attributed to the formation of the AF-predicted inter-molecular zinc finger between HBx and Spindlin1. Indeed, no other Cys residues are present in HBx_56-76_ sequence; Spindlin1 has not been reported to bind Zn^2+^ on its own.

**Figure S9.**
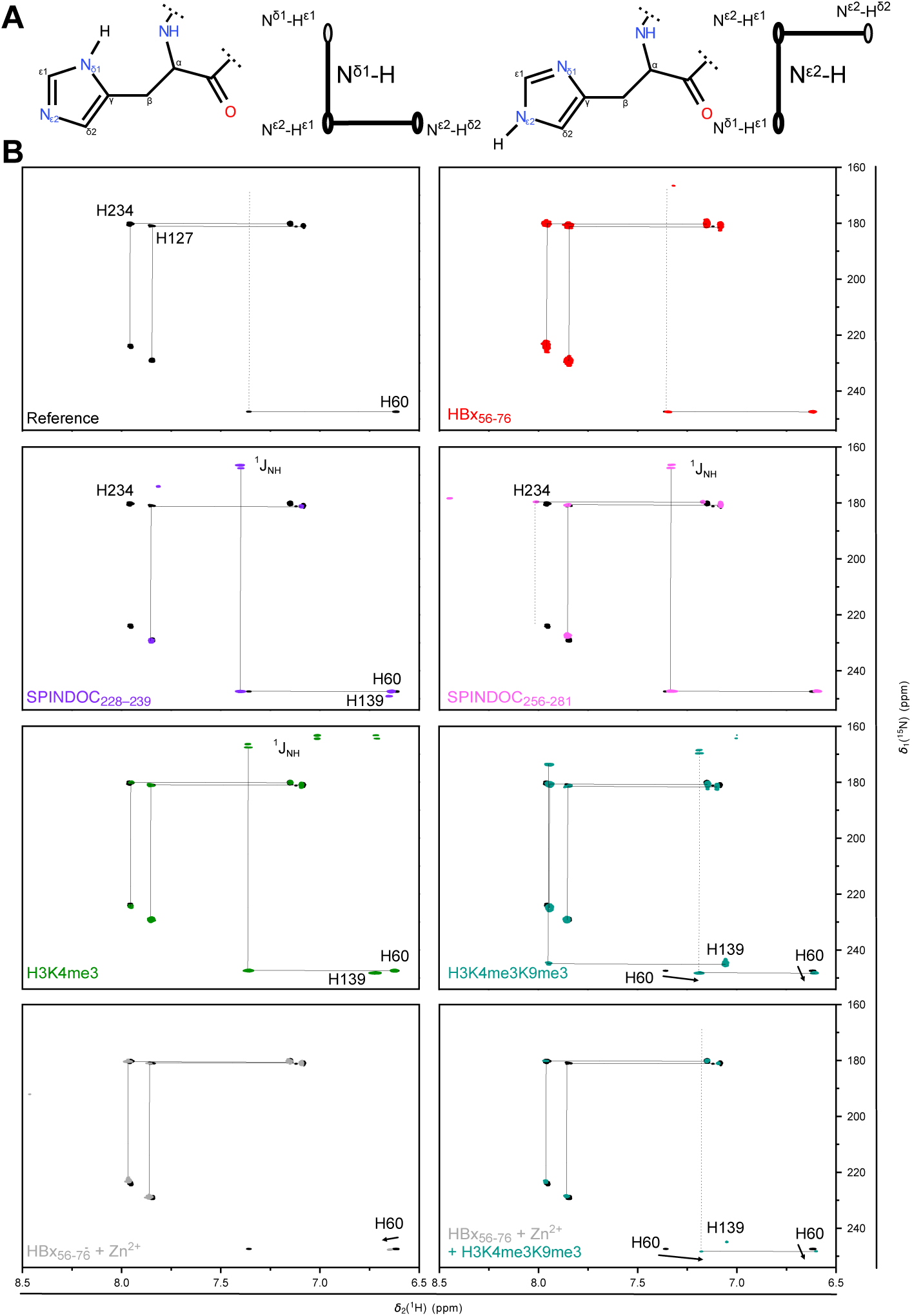
His residues on Spindlin1 report on binding peptides from HBx, SPINDOC and methylated histone tails. **A.** His side chains can exist in two tautomeric states, defined by which nitrogen atom in the imidazole ring is protonated: either the N^δ1^–H or N^ε2^–H tautomer. The equilibrium depends on pH, hydrogen-bonding environment, and metal coordination. Binding a partner protein can change sidechain conformation or solvent exposure and hence the tautomeric state. The associated ¹H–¹⁵N correlations appear at distinct chemical shift positions^66^. The resulting signals patterns are shown as schematics, from which the tautomers can be readily identified. **B.** Overlay of histidine sidechain correlations of free Spindlin1 (black contours, with sequential assignments) and in complex with peptides (in colour). For peptide-complexed Spindlin1, signals that experience CSPs upon binding are marked with arrows. The spectra were recorded on the same samples as the backbone amide correlation spectra in Fig. S8. Signals were assigned by deduction based on the observed CSPs and the position of His residues with respect to the peptide binding sites (Fig. S7C). The ^1^H-^15^N H(C)-N HMQC^67^ experiment was optimized for the detection of His sidechain correlations (^2^*J*_HN_ set to 45.45 Hz). The appearance of ^1^*J*_HN_ splittings in the indirect dimension, which is not ^1^H decoupled, implies protection of the N-H moiety from exchange. The signal of H252, which mediates the zinc finger, is detectable neither in the free nor in the bound state, as confirmed by mutagenesis. Addition of Zn^2+^ induces a CSP to H60, consistent with an effect of the zinc finger on the K9me3 pocket of Tudor 1 domain. Unlike full HBx_1-120_ protein (Fig. 5E), HBx_56-76_ peptide is outcompeted H3K4me3K9me3 even in the presence of Zn^2+^ the N-terminal hairpin of HBx is indispensable for high-affinity binding.

**Figure S10.**
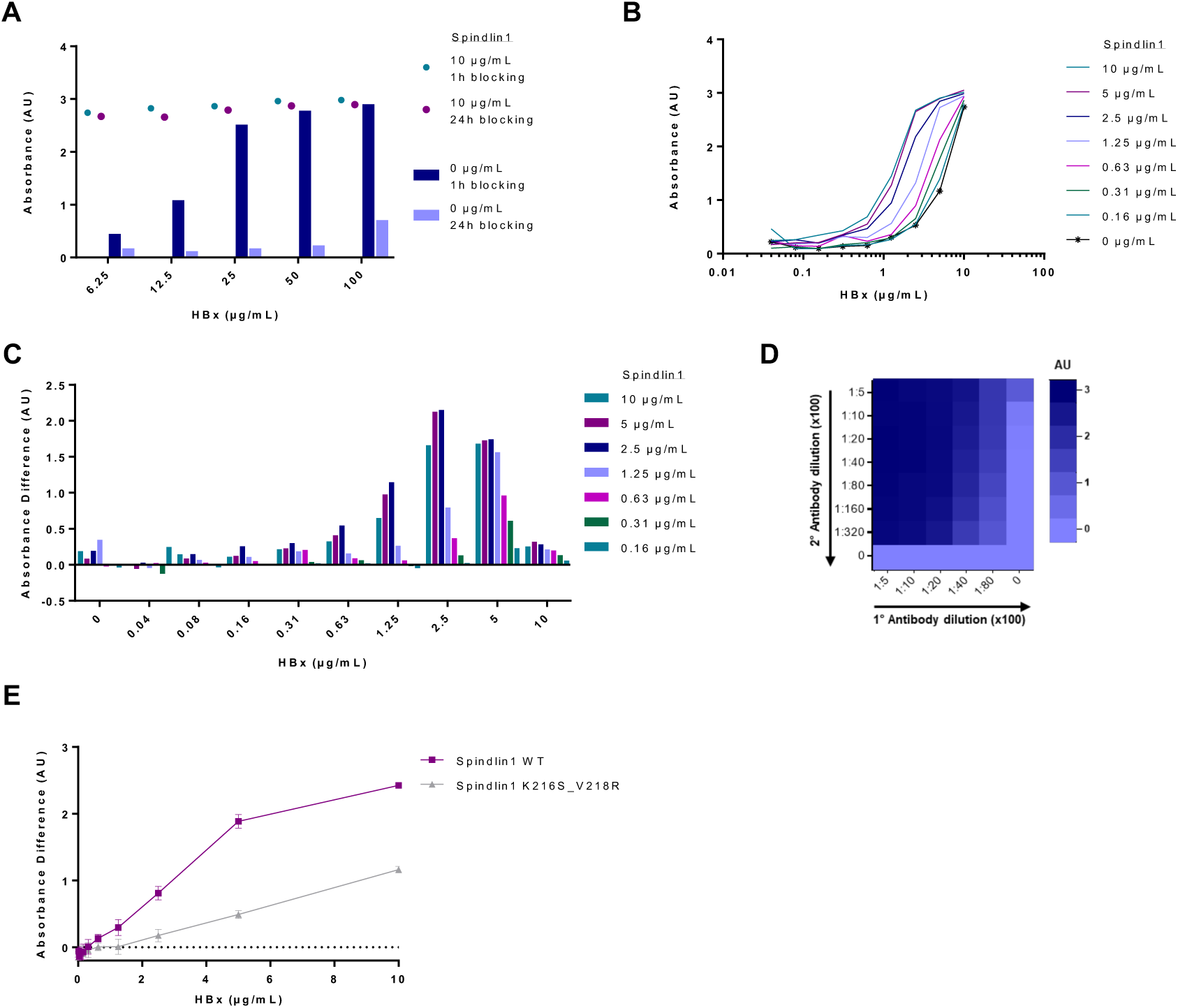
Optimization of ELISA setup for recombinant HBx_1-120_. A. Plates were coated with 0 µg/mL or 10 µg/mL Spindlin1 and blocked with PBS-Tm blocking buffer for 1h or overnight. Subsequently, Spindlin1 was detected using various amounts of recombinant HBx_1-120_ from 0–100 µg/mL. Unspecific HBx_1-120_-binding to the plate (no Spindlin1 coated) increases with HBx_1-120_ concentration, whereas it decreases with longer blocking time (blue and violet bars). On the contrary, the signal of Spindlin1 recognition is nearly unaffected by either HBx_1-120_ concentration or blocking time, implying a saturation of signal at the used protein amounts. B. To determine the optimal protein concentrations, plates were coated with serial dilutions of Spindlin1 from 0–10 µg/mL and blocked, before detection with a dilution series of 0–10 µg/mL HBx_1-120_. Shown are the uncorrected absorbance values in dependence of the used HBx_1-120_ concentration. Background signal of unspecific HBx_1-120_ binding to the plate (0 µg/mL Spindlin1) is shown with black asterisks. C. Background-corrected detection of Spindlin1 by recombinant HBx_1-120_. The signal drop at 10 µg/mL HBx_1-120_ results from unspecific binding to the ELISA plate, implying concentrations of 2.5 or 5 µg/mL HBx_1-120_ are sufficient for detection without relevant background signal. Similarly, Spindlin1 coated at 2.5 or 5 µg/mL (dark blue- and plum-coloured bars) result in good signal. D. Optimization of antibody concentration by serial dilution of 1° rabbit-anti-HBx and 2° anti-rabbit HRP-coupled antibodies. Relative absorbance signal (compared to lowest dilution of either 1° or 2° antibody) is above 90% for 1° antibody dilutions up to 1:2,000 and 2° antibody dilutions up to 1:8,000. Some unspecific binding of 2° antibody can be seen at dilutions of 1:500 and 1:1,000 but this is neglectable (<5% of relative signal) at higher dilutions. E. To verify specificity of HBx_1-120_ binding Spindlin1, Spindlin1_K216S/V218R_ harbouring two mutations at the β-hairpin site was used and shows reduction of HBx_1-120_ binding, in line with NMR observations.

## Methods

### Production of recombinant proteins

All chemicals were purchased from Sigma/Merck (Darmstadt, Germany), Carl Roth (Karlsruhe, Germany), SERVA (Heidelberg, Germany) unless otherwise stated.

#### HBx_1-120_

For NMR experiments, HBx_1-120_ protein genotype D (Uniprot P03165, natural variant R26C) was produced tag-free as detailed previously^26^. In summary, HBx_1-120_ was expressed in *E. coli* BL21(DE3) and recovered from inclusion bodies via solubilization in 8 M urea. The protein was purified in two steps via ion exchange and size exclusion chromatography in 6 M urea. The following point mutations were introduced using Q5 site-directed mutagenesis kit (New England Biolabs, Ipswich, USA): V12I, V21I, L58M, L71M L108G, V60G, C61A - C69A, L116M.

To obtain ^15^N, I-δ_1_-[^13^CH_3_], V/L-γ_2_/δ_2_*(proS)*-[^13^CH_3_, ^12^CD_3_], Mε-[^13^CH_3_] (referred to as ILVM-^13^CH_3_-) methyl-labelled HBx_1-120_, the previously published expression protocol^26^ was slightly modified. Cells were grown from a single colony in LB medium for 8 h at 37°C. The preculture was transferred into D₂O-based M9 minimal medium supplemented with ^15^NH₄Cl (1 g/L) and [^2^D, ^12^C]-glucose (3 g/L) as sole nitrogen and carbon sources to achieve an initial OD₆₀₀ ∼0.05. The preculture was incubated overnight at 37°C for minimal medium adaptation and then transferred to the final expression volume at an initial OD₆₀₀ ∼0.1 and grown at 37°C to OD₆₀₀ ∼0.6. At this point, DLAM-LVproS-^13^CH₃ Methyl Labeling Kit (NMR-Bio, L’Albenc, France) and L-methionine-(ε-^13^CH₃) (100 mg/L) (CORTECNET, Les Ulis, France) were added. After 40 min, α-ketobutyrate (L-isoleucine-δ₂-^13^CH₃ precursors) (60 mg/L) was added, followed 20 min later by induction with 1 mM isopropyl β-D-1-thiogalactopyranoside (IPTG). Cells were harvested after 4 hours of expression at 37°C by centrifugation, and HBx_1-120_ was purified as described previously^26^. Production of ^15^N labelled HBx_1-120_ was conducted similarly without the addition of precursors/amino acids in water-based M9 medium.

#### Bcl-xL

The expression construct encoded residues 1–44 and 85–212 of human Bcl-xL (Uniprot Q07817), lacking the transmembrane domain and the loop spanning the residues 45–84. This soluble domain is referred to as Bcl-xL here^68^. Bcl-xL was expressed and purified using the same protocol as that described for Spindlin1^26^. The hexa-histidine tag of Bcl-xL was cleaved using thrombin (Cytiva, Marlborough, MA, USA) overnight at room temperature and the cleaved protein was recovered by collecting the flow-through from a reverse IMAC (Immobilized metal affinity chromatography) using as running buffer 50 mM Tris-HCl pH 8.0, 300 mM NaCl, 40 mM imidazole and subsequent size exclusion. The ^15^N, I-δ_1_-[^13^CH_3_], V/L-γ_2_/δ_2_*(proS)*-[^13^CH_3_, ^12^CD_3_], Mε-[^13^CH_3_] (referred to as ILVM-^13^CH_3_-) labelled Bcl-xl was expressed as described for HBx_1-120_ except that the culture was incubated overnight at 32°C post IPTG induction^64^.

#### Spindlin1

The expression construct encoded residues 50–262 of human Spindlin1 (Uniprot Q9Y657), lacking the unstructured N-terminal segment of the protein with the addition of an N-terminal hexa-histidine tag and TEV (tobacco etch virus) protease cleavage site. The protein was expressed and purified as detailed previously^26^. The tagged protein was dialyzed overnight at room temperature against the standard buffer A (150 mM NaCl, 50 mM HEPES pH 7.4) in the presence of TEV. The cleaved protein was recovered by reverse IMAC as for Bcl-xL. The flow through was collected, concentrated and loaded onto a size exclusion HiLoad 16/600 Superdex 200 pg column (Cytiva, Marlborough, MA, USA) equilibrated with buffer A. ^15^N-labelled Spindlin1 was expressed as described previsouly^26^ with the addition of a medium adaptation step as described above for HBx_1-120_. The following point mutation combinations were introduced using site-directed mutagenesis: K216S-V218R, C96A-H252F.

### Peptide preparation

All peptides were synthetised by Genscript (Piscataway, USA). The sequences are listed in Table S1. SPINDOC_228-239,_ Histone H3K4me3_1-20,_ Histone H3K4me3K9me3_1-20_ peptides were solubilized in buffer A. SPINDOC_256–281_ and HBx_56-76_ peptides were only fully soluble in water and partially aggregated once mixed with buffer A to obtain NMR samples. Peptide concentrations were obtained by absorption measurement at 205 nm (NanoDrop One, ThermoFisher Scientific) by taking a series of peptide dilutions with water to mitigate the interference of HEPES at 205 nm. The extinction coefficients at 205 nm were determined based on the peptide sequence using the BESTSEL server^69^.

**Table S1.**
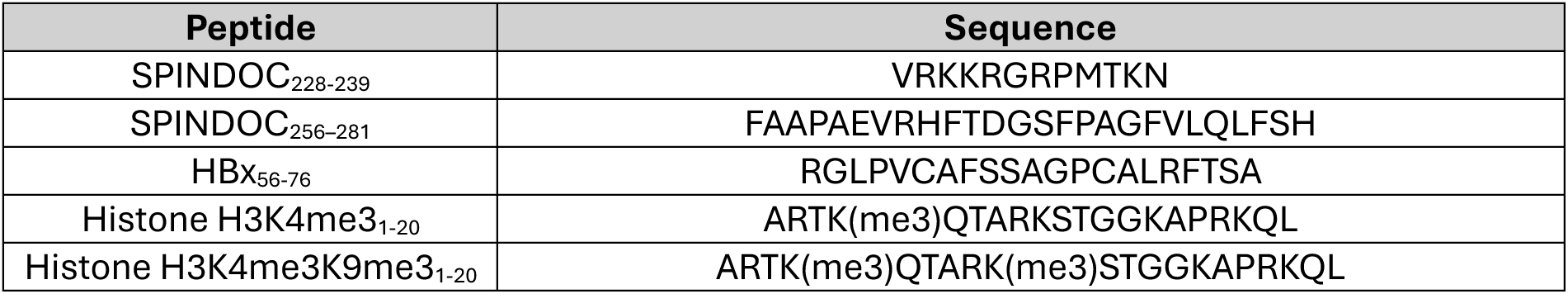
Peptide sequences used for NMR experiments. HBx_56-76_ was defined based on the NMR binding assay and the AF prediction as the minimal sequence required to mask the K4me3 and K9me3 binding pockets. SPINDOC_228-239_, also known as DOCPEP2, was previously identified to bind Spindlin1 and mask the K4me3 binding pocket. SPINDOC_256–281_, also known as DOCPEP3, corresponds to the SPINDOC segment docking as a β-hairpin onto the Tudor 3 domain of Spindlin1^42^.

### NMR sample preparation

For each NMR experiment involving HBx_1-120_, the protein was first dialysed from size exclusion buffer with 6 M urea (50 mM NaCl, 50 mM HEPES pH 7.4, 5mM DTT) to the buffer with 4 M urea (150 mM NaCl, 50 mM HEPES pH 7.4, 1 M L-arginine, 5mM DTT) and in a second dialysis step to a similar buffer containing 1.5 M urea. In the latter buffer, HBx_1-120_ was concentrated to 400 µM. The protein was then diluted eightfold: either with buffer A alone to obtain reference samples of free HBx_1-120_, or with host protein(s) or peptide in buffer A/water to obtain bound HBx_1-120_. Hence, samples were obtained of HBx_1-120_, either alone or in complexes, at a concentration of 50 µM in the final buffer (190 mM urea, 150 mM NaCl, 50 mM HEPES pH 7.4, 125 mM L-arginine). The DOSY reference samples of Bcl-xL or Spindlin1 alone were prepared at 50 µM in the same buffer.

Addition of Zn^2+^ was conducted by placing the NMR sample in a dialysis unit (6 kDa cutoff, Merck, Darmstadt, Germany) for at least 8 hours at 4°C against the NMR buffer supplemented with 200 µM ZnCl_2_. Addition of EDTA was conducted by spiking the sample with a 250 mM stock to obtain 1 mM EDTA final concentration.

For Spindlin1-detected experiments, the protein was concentrated in buffer A and mixed with the solubilized peptides. Addition of Zn^2+^ to those samples was conducted by placing the NMR sample in a dialysis unit (1 kDa cutoff, Merck, Darmstadt, Germany) for at least 6 hours at room temperature against buffer A supplemented with 400 µM ZnCl_2_.

### NMR spectroscopy

Solution-state NMR experiments were conducted on an Avance III Bruker spectrometer equipped with a TCI cryo probe at a magnetic field strength corresponding to ^1^H resonance frequency of 800 MHz. Sample temperatures were 4 °C for all experiments involving HBx_1-120_ and for the DOSY. Spindlin1-detected experiments (backbone amide and histidine sidechain correlations) were recorded at 37°C. Spindlin1 histidine sidechain correlation spectra were acquired using a SOFAST H(C)-N HMQC sequence from NMRlib^67^. HBx_1-120_ and Spindlin1 backbone amide correlation spectra were acquired using a standard SOFAST ^1^H-^15^N HMQC sequence from the Bruker library.

Translational diffusion coefficients were measured by recording pseudo-2D ^13^C-edited DOSY NMR spectra using a pulse scheme with a ^13^C-edited bipolar-gradient-pulse pair longitudinal eddy current delay (BPP-LED)^70^, similar to a ^15^N-edited BPP-LED experiment reported before^71^, additionally including presaturation for water suppression. Diffusion times Δ and gradient lengths δ were selected accordingly (Δ = 0.354 s, δ = 3.0 ms) and 16 gradient steps with 5-95/98% of the gradient intensity were recorded for each sample. For the experiments with Spindlin1 and Bcl-xL alone without ^13^C labelling, a pulse scheme with a stimulated echo, bipolar gradients and water suppression (^13^C-decoupled stebpgp1s19) was employed. Diffusion coefficients were determined by fitting the integrated intensities of the methyl groups and the gradient strengths to the Stejskal-Tanner equation. Not all 16 points were used for every fit.

Experimental parameters are listed in Supplementary information 2. All spectra were processed using TopSpin (Bruker; v 3.5 and 3.7) and analysed using CcpNmr AnalysisAssign (3.3.2.3)^72^.

### NMR Assignments

#### HBx_1-120_

The sequential resonance assignments of backbone amide correlations of HBx_1-120_ in presence of 1 M and of 190 mM urea were previously performed^27^ (Biological Magnetic Resonance Bank accession number 53376). The assignment covers all detectable residues 4–120 as well as all visible sidechain correlations in ^1^H-^15^N HMQC spectra. Methyl group correlations of interest in HBx_1-120_ were assigned by mutagenesis for residues V15, V21, L58, V60, L116, while M79, M103, L100, L108 were deduced from signal loss or CSPs observed upon binding either Bcl-xL (effects in C-terminal region) or Spindlin1 (effects in N-terminal region or residues 50–80).

#### Bcl-xL assignment

The assignment of N, I-δ_1_-[^13^CH_3_], V/L-γ_2_/δ_2_*(proS)*-[^13^CH_3_, ^12^CD_3_], Mε-[^13^CH_3_] (referred to as ILVM-^13^CH_3_-) labelled Bcl-xL was previously performed by Mizukoshi *et al.*^64^ for the same construct at 25°C. Every assigned signal could be transferred to our buffer condition at 4°C.

#### Spindlin1 assignment

The side chain correlations of Trp residues in Spindlin1, which are visible in the ^1^H-^15^N HMQC spectrum were assigned using the point mutants W151A, W165F, W62F, W72F in their free and, where necessary, peptide-bound states (with H3K4me3 and H3K4me3K9me3).

### Intensity loss evaluation and visualization

The intensity plots express the ratio of the peak intensities (based on heights) between the bound and free forms of HBx_1-120_ or Bcl-xL. The intensity plots of the HBx_1-120_-Bcl-xL complex in Fig. 2A originated from spectra recorded on HBx_1-120_ with ^15^N label, in contrast to the spectrum of ILVM- ^13^CH_3_-labeled HBx_1-120_ shown in Fig. S2A. All the other intensity plot are directly derived from the spectrum shown alongside in this manuscript.

The HBx_1-120_ intensity plot covers the region 4–120, as the first three residues of the sequence are not detected in our reference spectrum. Signals for residues 7, 12, 65, 98 have been removed from the analysis and are marked with ‘*’ as signal overlap impedes the peak height estimation. In case of the signals for residues 22–32, some of which show peak doubling in complex with Spindlin1, the peak height of the bound state was calculated as the sum of the two peaks. To calculate the error of the peak height, the spectral noise level was estimated by the built-in CcpNMR function.

The intensity loss was visualized in the AF models by applying a colour gradient from 0 to 1. The first three undetected residues were given the same intensity ratio as residue 4. Pro residues and others removed from the analysis were visualized with the mean value of the ratios of their neighbouring residues.

### Predictions of diffusion coefficients

Structure-based diffusion coefficients were obtained with HYDROPRO^32^. Calculations used an atomic-level primary model and shell calculation, hydrodynamic radius AER of 2.9 Å, six values of the radius of the minibeads, a partial specific volume of 0.73 cm^3^/kg, and an automatic choice of scattering data and number of intervals for the distance distribution. Calculations were run at 4°C, at a solvent density of 1.002 g/cm^3^ and a viscosity of 0.01858 P. The latter was determined by measuring the viscosity of the NMR buffer with an Anton Paar MCR 100 cone-plate rheometer. At 4 °C, the viscosity of the NMR buffer was measured at η = 1.858 mPa·s after recalibration based on water reference. The HYDROPRO predictions of 3-1-2 versus 3-2-1 topology (Fig. S3 and S4) from AF yielded identical *D* values.

Dudas et al.^33^ proposed an empirical approach to calculate the diffusion coefficient D of a folded and an intrinsically disordered protein at 287 K in water. They obtained two functions

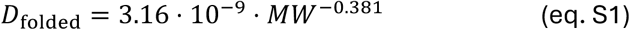

and

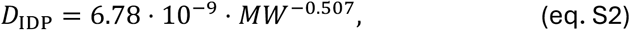

with the molecular weight MW (in g/mol), yielding the diffusion coefficient D (in m^2^/s). Based on our own viscosity measurement, the D obtained from those equations can be transposed to our buffer and temperature conditions following the equation

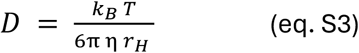

with k^B^ the Boltzmann constant, T the absolute temperature, η the viscosity and r^H^ the hydrodynamic radius. Thus, we obtain the empirical and recalibrated D coefficients for the three constructs used herein (Table S2).

**Table S2.** Empirically predicted diffsusion coefficients for isolated proteins. Values were calculated from the empirical Dudas model (eq’s. S1 and S2) and calibrated to the actual experimental conditions (eq. S3).

| Protein | $D_{\text{Dudas}}$ at 287 K in water<br>[10<sup>-11</sup> m<sup>2</sup>/s] | $D_{\text{Dudas}}$ at 277 K in buffer<br>[10<sup>-11</sup> m<sup>2</sup>/s] |
| --- | --- | --- |
| HBx<sub>1-120</sub> | 5.58 (IDP) | 3.41 |
| Bcl-xL | 7.26 (folded) | 4.43 |
| Spindlin1 | 6.72 (folded) | 4.10 |

### AlphaFold models

All AlphaFold models were obtained from AlphaFold server running AlphaFold 3^30^. The Bcl-xL and Spindlin1 sequences corresponded to the NMR constructs (residues 1–44/85–212 and 50–262, respectively), rather than the full-length proteins, with no cleavage scars included. In addition, Bcl-xL models included the three N-terminal residues left over after the thrombin cleavage. The models represented in this manuscript for the HBx_1-120_-Spindlin1 complex were not necessarily the best-ranking score but were selected to represent the best fit to the experimental data. Fig. S3 and S4 show the two models with the most extended interaction surface of residues HBx_50-80_ towards Spindlin1, which furthermore samples two alternative topologies of contacting the Tudor domain pockets (3-1-2 versus 3-2-1). Confidence scores of all the models in this manuscript are shown in Table S3.

When a Zn^2+^ ion was included in the computational model of the HBx_1-120_-Spindlin1 complex, we observed a slight reduction in the diversity of AF hits, suggesting that Zn^2+^ coordination promotes convergence among predicted structures. When all models were aligned on Spindlin1, the mean RMSD of the interacting HBx_50-80_ region relative to the average atomic coordinates decreased from 9.25 ± 1.67 Å without Zn^2+^ to 8.03 ± 1.04 Å when providing Zn^2+^ (N = 9 and 8, respectively); *t*(15) = 1.78, *p* = 0.099 (n.s.).

**Table S3.** AlphaFold 3 confidence metrics for predictions in this study. pTM: predicted template modelling score; ipTM: interface predicted template modelling score. Low confidence is obtained for HBx structures due to the large degree of disorder.

| Components | Global pTM score | ipTM score | Chain pTM (HBx; partner) | Figure |
| --- | --- | --- | --- | --- |
| HBx<sub>1-154</sub>, Zn<sup>2+</sup> | 0.25 | 0.84 | 0.25; - | Fig. 1B |
| HBx<sub>1-120</sub>, Zn<sup>2+</sup> | 0.20 | 0.63 | 0.20; - | Fig. 1C |
| HBx<sub>1-120</sub>, Bcl-xL | 0.56 | 0.20 | 0.16; 0.83 | Fig. 2B,D; S2B |
| HBx<sub>1-120</sub>, Spindlin1, topology 3-1-2 | 0.68 | 0.79 | 0.24; 0.84 | Fig. 3C,E; S3C |
| HBx<sub>1-120</sub>, Spindlin1, Zn<sup>2+</sup>, topology 3-1-2 | 0.69 | 0.81 | 0.26; 0.85 | Fig. 4B, 5C, S4D, S7A |
| HBx<sub>1-120</sub>, Spindlin1, topology 3-2-1 | 0.68 | 0.79 | 0.24; 0.85 | Fig. S3C |
| HBx<sub>1-120</sub>, Spindlin1, Zn<sup>2+</sup>, topology 3-2-1 | 0.70 | 0.81 | 0.27; 0.85 | Fig. S4D |

### ELISA assay

ELISA plates (Nunc MaxiSorp® flat-bottom) were coated with 100 µL/well of recombinant Spindlin1 at various concentrations in coating buffer (50 mM carbonate, pH 8) and incubated with slight agitation at 4°C over-night. The next day, plates were washed three times with 200 µL/well PBS-T (PBS + 0.05% Tween-20) and blocked with 100 µL/well PBS-Tm (5% milk in PBS-T) for various time intervals. All subsequent sample incubation steps were performed with 100 µL/well at room temperature with slight agitation and washing with three times 200 µL/well PBS-T in between. Protein concentrations were determined by NanoDrop, Bradford assay^73^ and/or SDS-PAGE and varied in the optimization process. HBx_1-120_ was stepwise dialysed and diluted as described above. Subsequently, dilution in PBS-Tm was performed to achieve a concentration of 2.5 µg/mL to 100 µg/mL. HBx_1-120_ was incubated on Spindlin1-coated plates for 1h. Detection of Spindlin1-bound HBx_1-120_ was performed using an in-house polyclonal rabbit-anti-WHBx_1-50_ (Woodchuck) antibody, followed by HRP-conjugated-anti-rabbit antibody (Thermo Fisher), both diluted in PBS-Tm. Inhibitors/peptides were co-incubated/co-diluted at various concentrations with HBx_1-120_. Detection was performed with 100 µL/well 3,3′,5,5′-Tetramethylbenzidine (TMB) substrate-solution (Merck, Sigma) and stopped after 20-30 minutes using 100 µL/well 1 M HCl. Absorbance was measured on a BMG plate reader at 450 nm (see also Fig. S10). For the data shown in Fig. 5F, recombinant Spindlin1 at 2.5 µg/mL was immobilized and detected using 5 µg/mL refolded HBx_1-120_. Primary HBx-specific and secondary HRP-coupled antibodies were used at dilutions of 1:2,000 and 1:8,000, respectively.

### Multiple sequence alignment

Sequences IDs used for the sequence alignment (Fig. S1A) are available in Supplementary Information 3. Alignment was done using Clustal Omega^74^ and visualized with JalView 2.11.5^75^. Statistical analysis of the cysteine abundance (Fig. 6B) in proteins belonging to HBx isoforms, viruses, or *Homo sapiens* was obtained based on extracting sequences from the UniProt database^76^ using the following queries: HBx sequences–“gene: x AND taxonomy_id:10407 AND reviewed:true”; human sequences–“taxonomy_id:9606 AND reviewed:true”; Virus sequences– “taxonomy_id:10239 AND reviewed:true”.

### Search for inter-molecular zinc fingers in the Protein Data Bank

Structures of inter-molecular zinc fingers between viral and eukaryotic proteins reported in the PDB were identified using a series of database queries. First, UniProt accessions of viral proteins (taxonomy ID 10239) featuring a PDB structure were obtained using the UniProt API with following query: “taxonomy_id:10239 AND database:pdb”. Corresponding PDB accessions were gathered using the “xref_pdb” field. PDB was queried for a list of proteins present in the corresponding structures, in the UniProt accession format, using PDBe’s mapping API. These UniProt accessions were filtered for eukaryotic proteins using “taxnonomy_id:2759”, allowing to identify PDB structures featuring both a viral and eukaryotic proteins, which yielded 2509 structures. For each entry, the field “nonpolymer_bound_components” was extracted using the RCSB API, and the result was filtered for entries containing Zn^2+^. mmCIF structure files for each of the resulting 305 entries were parsed to identify all protein chains present within a 3 Å radius around each Zn^2+^, then automatically filtered for ions that are simultaneously coordinated by a chain from a eukaryotic and a viral protein, respectively. The resulting 11 structures were inspected manually, resulting in six PDB entries, all featuring the complex of HIV-1 tat and the human CDK9/cyclin T1 complex (PDB-IDs: 3MI9, 3MIA, 4OR5, 6CYT, 5L1Z, 4OGR). Indeed, the interface between cyclin T1 and HIV-1 tat is stabilized by a shared Zn^2+^, with the viral tat contributing three of the four coordinating Cys residues. Notably, the corresponding regions of tat is Cys-rich, suggesting a similar interaction strategy to that employed by HBx.

### Visualization

Molecular graphics and analyses were performed with Chimera version X (1.10)^77^.

### Data availability

Data supporting the findings of this work are available within the Article, the Methods, the Supplementary Figures and Information. Raw data from NMR data and from *in-silico* modelling are available from the corresponding author upon request.

## Acknowledgements

The authors thank Prof. Björn Burmann (University of Gothenburg) for sharing the BPP-LED pulse sequence, Prof. Wolfgang Friess and Carolin Thilmann (Ludwig Maximilian University of Munich) for the viscosity measurements, Prof. Franz Hagn (Technical University of Munich) for the Bcl-xL plasmid, and Christa Kuhn (University of Heidelberg) for providing the WHBx antibody.

## Supplementary information 1

**Table S4.** Collection of affinities previously determined by ITC for peptides derived from HBx, SPINDOC or H3 histones towards Bcl-xL or Spindlin1. In this study, the HBx_61-80_ peptide was replaced by HBx_56-76_ as the latter fully covers the AF-predicted interaction region between Spindlin1 and HBx. The BH3-like helix in HBx is truncated in the HBx_1-120_ isoform. As a result, the helix shifts from residues 101–136 to 101–120, concomitant with a loss of affinity for Bcl-xL.

| Peptide | Affinity for Spindlin1 | Affinity for Bcl-xL |
| --- | --- | --- |
| HBx<sub>2-21</sub> | 1.1 $\mu\text{M}$<sup>24</sup> | |
| HBx<sub>61-80</sub> | 60.5 $\mu\text{M}$<sup>24</sup> | |
| HBx<sub>101-136</sub> | | 89 $\mu\text{M}$<sup>31</sup> |
| HBx<sub>101-120</sub> | | 222 $\mu\text{M}$–356 $\mu\text{M}$<sup>31</sup> |
| SPINDOC<sub>226-239</sub> | 15.4 $\mu\text{M}$<sup>42</sup> | |
| SPINDOC<sub>256-281</sub> | 30 nM<sup>42</sup> |  |
| H3K4me3<sub>1-20</sub> | 57.2 $\mu\text{M}$<sup>42</sup> | |
| H3K4me3K9me3<sub>1-20</sub> | 7.6 nM<sup>42</sup> |  |

## Supplementary information 2

Supplementary information 2 Experimental parameters for NMR measurements
| Experiment # | Experiment | Points | NS | Spectral widths (ppm) | Acquisition times (ms) | Carrier frequencies (ppm) |
| --- | --- | --- | --- | --- | --- | --- |
| 1 | sfhmqcf3gp-ph (Bruker library) | 2048-512 (H-N) | 128 | 16.02-32 (H-N) | 79.87-98.63 (H-N) | 4.71-101 (H-N) |
| 2 | sfhmqcf3gp-ph (Bruker library) | 2048-512 (H-N) | 256 | 16.02-32 (H-N) | 79.87-98.63 (H-N) | 4.71-101 (H-N) |
| 3 | BB sofast HMQC2 | 1536-256 (H-C) | 64 | 12.98-25 (H-C) | 73.93 – 25.44 (H-C) | 4.71-20 (H-C) |
| 4 | BB sofast HMQC2 | 1536-512 (H-C) | 128 | 12.98-25 (H-C) | 73.93 – 50.88 (H-C) | 4.71-20 (H-C) |
| 5 | sfhmqcf3gp-ph (Bruker library) | 1536-512 (H-C) | 128 | 12.98-25 (H-C) | 73.93 – 50.88 (H-C) | 4.71-20 (H-C) |
| 6 | sfhmqcf3gp-ph (Bruker library) | 1536-512 (H-C) | 32 | 12.98-25 (H-C) | 73.93 – 50.88 (H-C) | 4.71-20 (H-C) |
| 7 | SOFAST H(C)-N HMQC for histidine sidechain | 1276-768 (H-N) | 128 | 12.06-105 (H-N) | 73.93 – 50.88 (H-N) | 4.72-202.5 (H-N) |

Buffers compositions for NMR samples
| Buffer # | Composition |
| --- | --- |
| 1 | 190 mM urea, 125 mM L-Arginine, 150 mM NaCl, 50 mM HEPES, pH 7.4 |
| 2 | 150 mM NaCl, 50 mM HEPES, pH 7.4 |

NMR samples used for NMR experiments
| Sample | Labelling scheme | Buffer | Protein conc. | NMR experiment # | Figures |
| --- | --- | --- | --- | --- | --- |
| HBx1-120-detected experiments |  |  |  |  |  |
| HBx1-120 reference | <sup>2</sup>D, <sup>15</sup>N, <sup>13</sup>C(ILVM)methyl-labelled | 1 | 50 μM | 1 | Fig. 1D |
| + Bcl-xL complex |  | 1 |  |  | Fig. S2A |
| + Spindlin1 complex |  | 1 |  |  | Fig. 3B,D; S3A; S4A,B; S5A |
| + Bcl-xL + Spindlin1 ternary complex |  | 1 |  |  | Fig. S3D |
| + Spindlin1 Zn<sup>2+</sup> complex |  | 1 + 200 μM Zn<sup>2+</sup> |  |  | Fig. 4D; S4A, B; S5A |
| + Spindlin1 Zn<sup>2+</sup> EDTA complex |  | 1 + 200 μM Zn<sup>2+</sup> + 1 mM EDTA |  |  | Fig. 4D; S4A,B |
| HBx1-120 C61A C69A reference + Spindlin1 C96A H252F complex + Zn<sup>2+</sup> | <sup>15</sup>N-labelled | 1 | 50 μM | 1 | Fig. S5A,B |
| HBx1-120 Spindlin1 complete H3K4me3K9me3 titration (reference, complex, complex + peptide, + Zn<sup>2+</sup>) |  | 1 |  |  | Fig. 5E |
| HBx1-120 Spindlin1 complete H3K4me3K9 titration (reference, complex, complex + peptide, + Zn<sup>2+</sup>) |  | 1 |  |  | Fig. S6C |
| HBx1-120 Spindlin1 complete SPINDOC<sub>228-239</sub> titration (reference, complex, complex + peptide, + Zn<sup>2+</sup>) |  | 1 |  |  | Fig. S6B |
| HBx1-120 reference | <sup>15</sup>N-labelled | 1 | 50 μM | 2 | Fig. 3F; S3B |
| HBx1-120 reference + Spindlin1 complex + SPINDOC<sub>256-281</sub> |  | 1 |  |  | Fig. 5E |
| HBx1-120 reference + Spindlin1 K216S_V218R complex |  | 1 |  |  | Fig. 3B |
| HBx1-120 reference | <sup>2</sup>D, <sup>15</sup>N, <sup>13</sup>C(ILVM)methyl-labelled | 1 | 50 μM | 3 | Fig. 3F; S3B |
| + Spindlin1 complex |  | 1 |  |  | Fig. 3F; S3B |
| + Spindlin1 Zn<sup>2+</sup> complex |  | 1 |  |  | Fig. 4E; S4C |
| Bcl-xL-detected experiments |  |  |  |  |  |
| Bcl-xL reference | <sup>2</sup>D, <sup>15</sup>N, <sup>13</sup>C(ILVM)methyl-labelled | 1 | 50 μM | 4 | Fig. 2B,D; S2C,D,E |
| + HBx1-120 |  | 1 |  |  | Fig. 2B,D; S2C,D,E |
| Spindlin1-detected experiments (Spindlin1 + peptides) |  |  |  |  |  |
| Spindlin1 reference | <sup>15</sup>N-labelled | 2 | 200 μM | 5, 7 | Fig. S8A,B,C; S9B |
| + HBX<sub>56-76</sub> |  | 2 |  | 5, 7 | Fig. S8C; S9B |
| + HBX<sub>56-76</sub> + Zn<sup>2+</sup> |  | 2 + 600 μM Zn<sup>2+</sup> |  | 5, 7 | Fig. S8C; S9B |
| + H3K4me3K9me3 |  | 2 |  | 5, 7 | Fig. S8B; S9B |
| + H3K4me3K9 |  | 2 |  | 5, 7 | Fig. S8B; S9B |
| + SPINDOC<sub>228-239</sub> |  | 2 |  | 6, 7 | Fig. S8A; S9B |
| + SPINDOC<sub>256-281</sub> |  | 2 |  | 6, 7 | Fig. S8A; S9B |

## Supplementary information 3

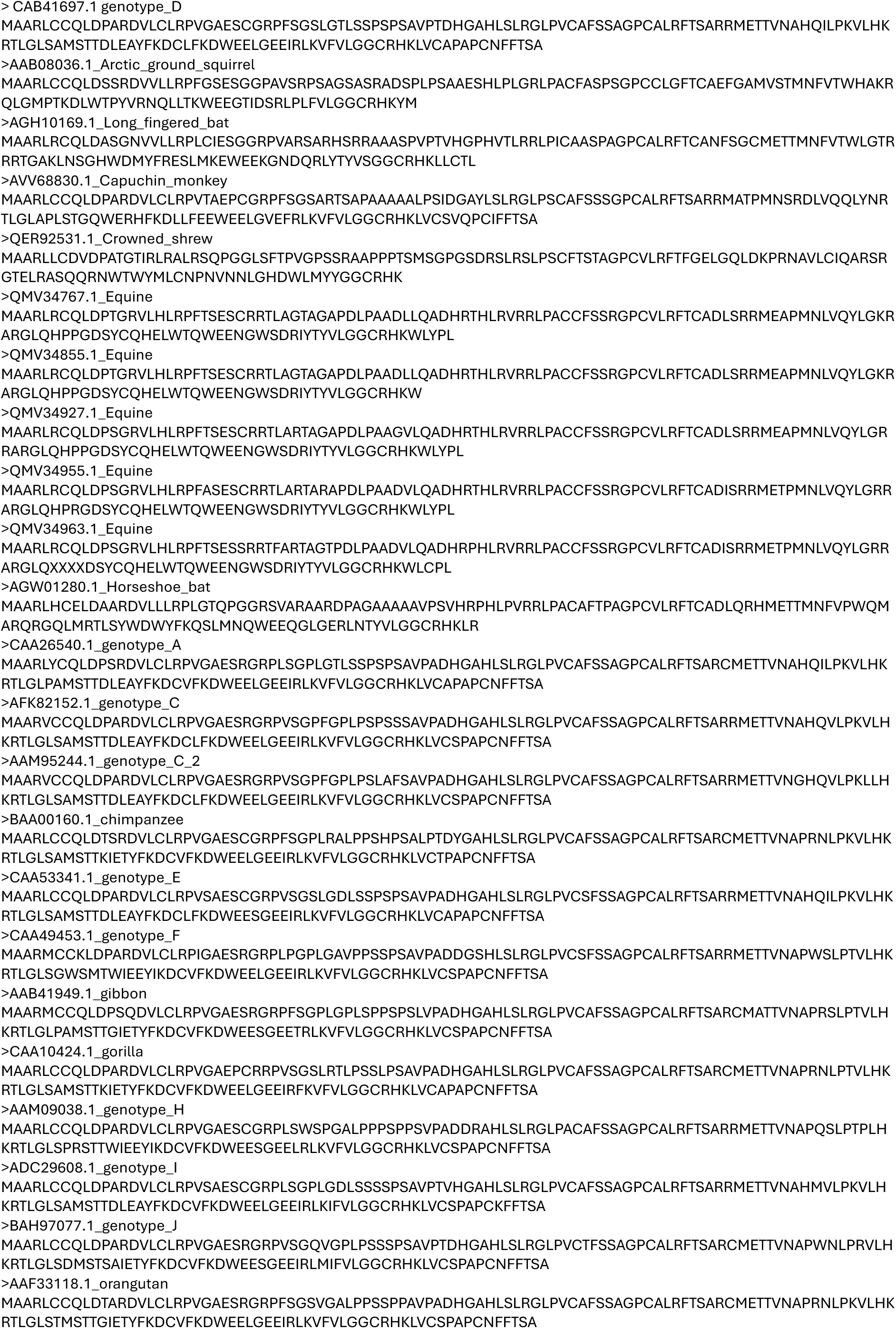
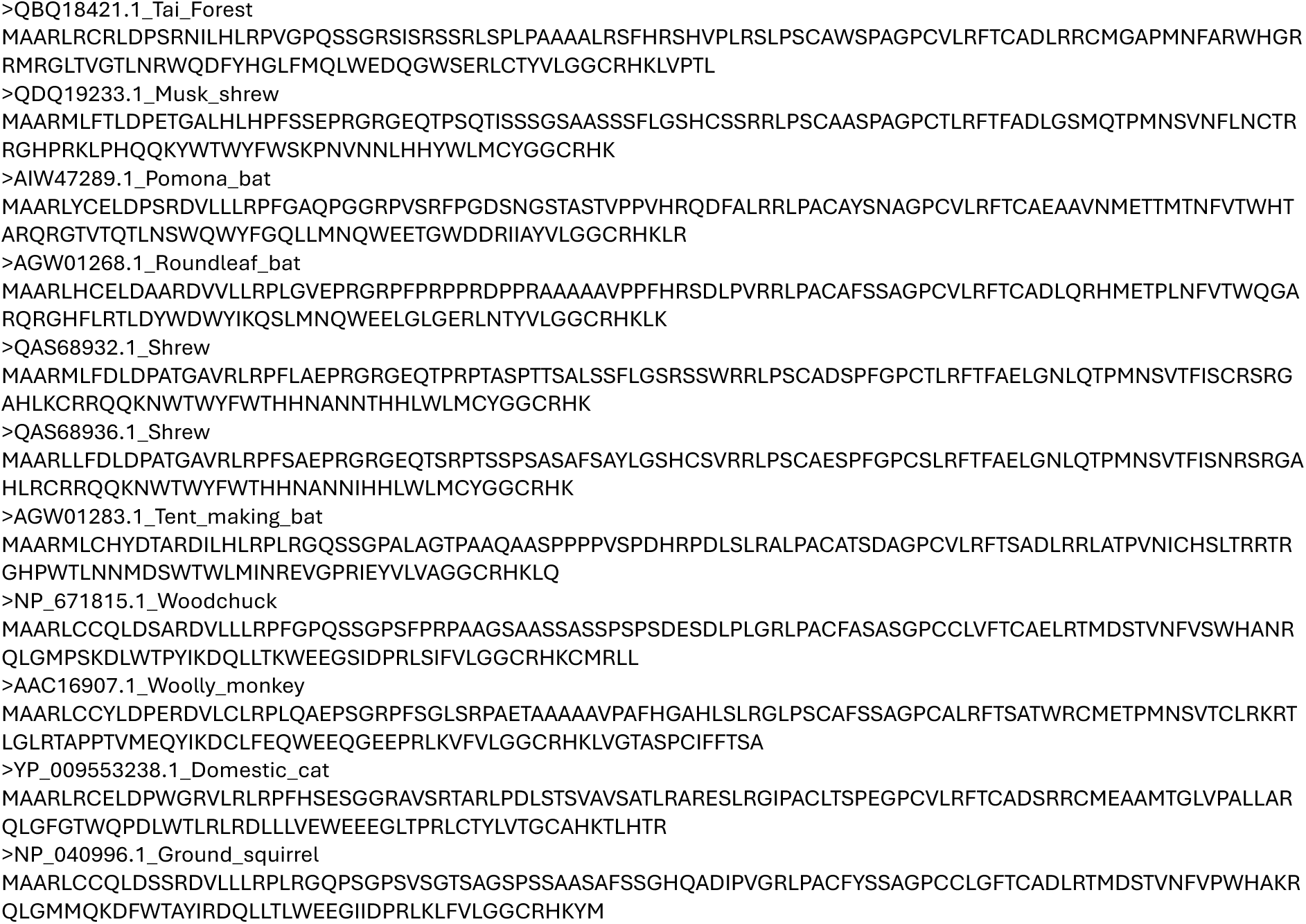

